# Integrative Genomic and Metabolomic Analysis of *Terribacillus aidigensis* KB25 Highlights Its Antifungal Activity against *Phytophthora infestans* and Adaptive Responses to Salt Stress

**DOI:** 10.64898/2026.02.07.704535

**Authors:** Gözde Koşarsoy Ağçeli, Ömür Baysal, Ragıp Soner Silme, Ümran Çapar

## Abstract

This study characterizes *Terribacillus aidigensis* strain KB25, a novel halotolerant isolate from a thermal spring, exhibiting potent antifungal activity against the late blight pathogen *Phytophthora infestans*. Whole-genome sequencing and electron microscopy revealed significant physiological adaptations to salt stress and a rich genomic repertoire encoding secondary metabolites. Metabolomic profiling of the bacteria-fungus interaction demonstrated upregulated cofactor biosynthesis and energy metabolism, specifically identifying antimicrobial terpene derivatives as key inhibitory agents. Complementary molecular docking studies provided mechanistic insights, predicting high-affinity binding between bacterial sporulenol and the *P. infestans* RxLR effector. Notably, the analysis indicated a strong structural interaction between the bacterial ABC-type proline/glycine betaine permease and the glutamate receptor, suggesting a mechanism for mediating plant stress tolerance. These findings validate the dual efficacy of KB25 which indicates direct pathogen suppression via bioactive metabolite secretion and the potential modulation of host stress signalling. Integrated genomic-metabolomic analysis was also proved our findings. Consequently, *T. aidigensis* KB25 represents a promising, sustainable biocontrol agent for managing late blight, particularly within saline agro-ecosystems.

**Graphical abstract:** 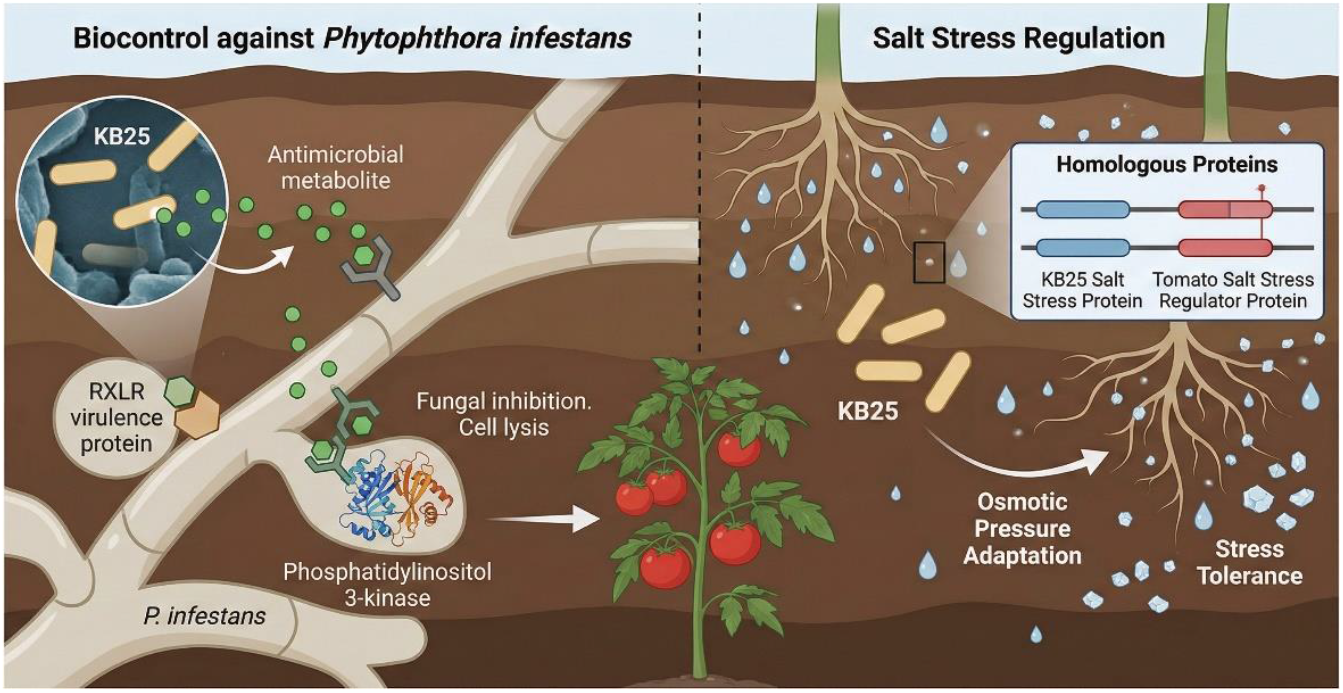

## INTRODUCTION

Climate change, population growth, and the depletion of freshwater resources have collectively intensified global food security challenges (Turral et al. 2011) Among the major environmental factors threatening agricultural productivity, soil salinization has emerged as one of the most critical abiotic stresses worldwide (Etesami and Glick 2020; Jiang et al. 2025). Excessive salt accumulation in the rhizosphere disrupts osmotic balance, reduces nutrient uptake and water potential. It causes ion toxicity and oxidative stress, leading to severe growth inhibition and yield loss (AbuQamar et al. 2024). Furthermore, salinity alters the composition and functionality of soil microbial communities, thereby weakening plant defense mechanisms and predisposing plants to opportunistic soil-borne pathogens such as *Fusarium, Rhizoctonia*, and *Phytophthora* species (Ha-Tran et al. 2021; Ould Ouali et al. 2024). The combined effects of salt stress and pathogen infection represent a major challenge to sustainable crop production, particularly in arid and semi-arid agroecosystems (Ullah et al. 2021; Sahu et al. 2023; Nachshon 2018).

*Phytophthora infestans*, the oomycete responsible for late blight in *Solanaceae*, employs a sophisticated hemibiotrophic lifestyle driven by a vast repertoire of effector proteins (Fry 2008; Park et al. 2026). Its pathogenicity hinges on the secretion of various RxLR effectors, which suppress host immunity and manipulate subcellular compartments like the nucleus and chloroplasts to facilitate colonization (Xu et al. 2024; Park et al. 2026). Recent genomic analyses reveal that *P. infestans* maintains high virulence plasticity through rapid effector evolution, allowing it to overcome resistant host genotypes and thrive in diverse agricultural environments (Coomber et al. 2024).

Soil salinity constitutes a critical abiotic constraint that significantly modulates the sensitivity of Solanaceae crops to late blight caused by various *Phytophthora* species. While physiological responses to osmotic stress prioritize ion homeostasis, this often compromises biotic resistance through signalling crosstalk. Specifically, the upregulation of abscisic acid (ABA) under salt stress frequently antagonizes salicylic acid (SA) pathways essential for immunity against hemibiotrophs. Consequently, salt-stressed hosts often exhibit reduced hypersensitive responses and compromised cell wall integrity, thereby facilitating pathogen ingress and colonization even in tolerant genotypes (Snapp et al, 1991; Preuett et al. 2016; Bai et al. 2018).

In this context, plant-microbe interactions have gained increasing attention as a pivotal component of plant stress physiology. Plants respond to environmental stress, not only through intrinsic metabolic and genetic mechanisms but also via symbiotic associations with microorganisms inhabiting the rhizosphere and endophytic compartments. Plant growth-promoting bacteria (PGPB) play a crucial role in mitigating both abiotic stresses (such as salinity, drought, heat, and heavy metals) and biotic challenges (including pathogens and pests) (Compant et al. 2005; Poria et al. 2022). These bacteria employ diverse mechanisms such as the synthesis of phytohormones (IAA, cytokinins, gibberellins), the production of ACC deaminase to lower ethylene stress, the secretion of siderophores and exopolysaccharides, and the modulation of antioxidant enzyme activity to enhance plant stress tolerance (Baysal et al. 2013; Backer et al. 2018; Egamberdieva et al. 2019). Additionally, PGPB can stimulate *induced systemic resistance* (ISR) by activating defense-related gene expression and modulating signal transduction pathways involving jasmonic acid and salicylic acid (Etesami and Glick 2020; Timmusk et al. 2017).

Recently, halophilic and halotolerant PGPB have been recognized as promising eco-biotechnological tools to enhance plant resilience under saline conditions (Meinzer et al. 2023; Reang et al. 2022). These microorganisms survive in hypersaline environments by synthesizing osmoprotectants, exopolysaccharides, compatible solutes, and bioactive secondary metabolites that not only protect bacterial cells but also improve soil aggregation, modulate ion transportation, and induce osmotic regulation in plants (Li et al. 2025a). Several halophilic strains have shown antifungal compounds production that suppress phytopathogens even under high-salt stress, which indicate their potential as dual-function bioinoculants since they are capable of promoting plant growth and protection against diseases (Pallavi et al. 2023; Ould Ouali et al. 2024).

Despite these advances, the mechanistic understanding of halophilic bacterial functions remains limited. Key knowledge gaps persist regarding the genetic and molecular basis of salt tolerance, antifungal metabolite biosynthesis, quorum sensing networks, and biofilm-mediated rhizosphere colonization (Jiang et al. 2025; Petrillo et al. 2021). The integration of multi-omics approaches including genomics, metabolomics and molecular simulations offers powerful tools to elucidate the regulatory networks that underpin halophilic bacterial adaptation and plant-beneficial activity. Such insights are crucial for advancing the rational design of next-generation bioinoculants that can function effectively under complex environmental stress conditions (Mashabela et al. 2025)

Within this framework in another study, *Terribacillus aidigensis*, a moderately halophilic bacterium frequently isolated from saline soils, has drawn increasing interest due to its ability to tolerate high salt concentrations and produce diverse enzymes and metabolites (Liu et al. 2010). However, its plant-beneficial properties, antifungal potential, and stress adaptation mechanisms remain poorly understood. To date, limited information on *Terribacillus* genus exists on its genomic determinants associated with osmotic stress tolerance, antimicrobial biosynthetic gene clusters (BGCs), and biofilm formation behaviour under saline conditions (Su et al. 2022; Petrillo et al. 2021).

In our study, the antifungal potential, biofilm formation capacity under different salinity levels, antibiotic susceptibility profile, and the genomic features of *T. aidigensis* isolated from a thermal spring in purpose of salt stress regulation and biocontrol efficiency against *P. infestans* were investigated. These analyses aimed to identify genes involved in osmotic adaptation, ion homeostasis, and secondary metabolite biosynthesis. By integrating phenotypic and genomic data enriched with metabolomics analysis provides novel insights into the biotechnological potential of *T. aidigensis* as a sustainable microbial resource for enhancing plant health and mitigating salinity stress in agricultural ecosystems.

## MATERIAL AND METHODS

### Sampling, isolation and characteristics of microorganism

Soil samples intended for microbial isolation were aseptically collected in sterile Falcon tubes from the outlet of a thermal spring located in the Ayvacık district of Çanakkale, Türkiye (39.57754° N, 26.16831° E). Water samples were inoculated onto Luria-Bertani Agar (Sigma) using the streak plate method to obtain single colonies. The inoculated plates were incubated at 50°C to promote the growth of thermophilic microorganisms. Among the isolates, the strain designated as KB25 (**K**oşarsoy-Ağçeli, **B**aysal, 20**25**) was selected for further analysis. This strain was preserved as a stock culture at −80°C for use in subsequent experimental studies.

Colony morphology was examined, and subsequently, Gram staining was performed. Biochemical characterization of KB25 colonies was performed using various culture media. Appropriate media were used for urease, indole, citrate, and other relevant tests. Bacterial morphology was visualized using a scanning electron microscope (SEM). Additionally, optimization of growth temperatures (25°C, 37°C, 45°C) and salt tolerance tests at various concentrations between 1% and 15% NaCl were tested.

### Biofilm Formation Potential at Different Salt Levels

The biofilm-forming potential of the microorganism was evaluated under different temperature and salt concentration conditions. The 24-hour microorganism culture was prepared at a turbidity of 1.0 McFarland standard (CLSI 2012). LB broth media containing 0-10% NaCl and the microbial cultures were added to the wells of microtiter plates. The plates were incubated at 25°C, 37°C, and 45°C for 24 hours. Following incubation, the wells were gently washed to remove non-adherent cells. The formed biofilms were stained with 0.01% crystal violet for 20 minutes. Unbound dye was removed by washing the wells three times with sterile distilled water. Subsequently, the bound dye was solubilized using 200 μL of ethanol, and the absorbance was measured at 570 nm to quantify biofilm formation.

### Scanning Electron Microscopy Observation

In order to prepare of SEM samples, the bacterial broth was first centrifuged to obtain a cell pellet, which was then washed three times with phosphate-buffered saline to remove residual medium components. The pellet was subsequently fixed by adding 0.25% glutaraldehyde prepared in sodium phosphate buffer (pH 7.2) and incubated at room temperature for 30 minutes, followed by overnight incubation to ensure complete fixation. After fixation, the cells were washed three additional times with sodium phosphate buffer and collected again by centrifugation. Dehydration of the samples was carried out using a graded ethanol series at concentrations of 30%, 50%, 70%, 80%, 90%, and 100%, with each step performed for 10 minutes to gradually remove water from the cells. The final dehydration step included incubation in 100% ethanol for 1 hour to ensure complete removal of moisture. Finally, SEM stubs were prepared by applying adhesive tape, onto which the dehydrated bacterial samples were mounted for subsequent imaging (Ali et al. 2021; Minuti et al. 2023).

### Antibiotic Susceptibility Testing

The antibiotic susceptibility test was conducted using the disk diffusion method, following the Clinical and Laboratory Standards Institute guidelines (CLSI 2021), with minor modifications to the medium employed (Erhonyota et al. 2023). The 24-hour microorganism culture was adjusted to a 0.5 McFarland turbidity (10^8^ cells/mL) and inoculated onto Mueller-Hinton agar containing 5% NaCl, with 100 µL of the culture. The antibiotic discs were placed on the plates, which were then incubated at 25°C, 37°C, and 45°C for 24 hours. The antibiotics used were Imipenem (IPM); Amikacin (AK); Polymyxin B (PM); Azithromycin (AZ); Cephalexin (CEP); Ceftriaxone (CRO); Cefoperazone (CFP); Metronidazole (MTZ); Furazolidone /Metronidazole (F/M); Netilmicin 30 µg (NET30); Fusidic Acid 10 µg (FA10); Amoxicillin-Clavulanate (AMC); Levofloxacin 5 µg (LEV5); Gentamicin 10 µg (CN10); Norfloxacin 5 µg (NB5); Cefixime (CFM); Cephalothin 30 µg (CE30); Moxifloxacin (MOX); Piperacillin-Tazobactam (TZP); Cefoperazone-Sulbactam 30 µg (ZOX30); Oxacillin 1 µg (OX1); Nitrofurantoin 300 µg (F300); Amikacin 30 µg (AN30); Ceftazidime 30 µg (CAZ30); Clindamycin 2 µg (DA2); Gentamicin (CN); Rifampin 5 µg (RA5); Cefoperazone-Sulbactam 105 µg (SCF105); Neomycin 10 µg (NN10); Cefazolin 30 µg (CZ30); Ofloxacin (OFX). After 24 hours, the inhibition zones were measured in millimetres. The experiment was performed with 3 replicates.

### Antagonistic interaction assay against *P. infestans*

*T. aidigensis* was incubated in Nutrient Broth (NB) at 28°C and 150 rpm for 24 hours. The performed culture was spread-seeded onto Nutrient Agar (NA) plates and incubated at 28°C for 24 hours. A thin layer of water agar was then spread on the culture surface and allowed to set. A layer of Potato Dextrose Agar (PDA) cooled to 45°C was then placed on the same plates. After the PDA layer solidified, a 5 mm x 5 mm block of *P. infestans* mycelium was placed in the centre. The plates were incubated at 27°C for 7 days, and fungal growth was visualised at 1, 3, 5, and 7 days after inoculation. As a control group, only *P. infestans* culture was used in PDA medium, and the experiment was performed in three replicates (Bozkurt et al. 2025).

### Salt stress regulation effect of KB25 on tomato

The *T. aidigensis* culture used in the experiment was propagated in NB medium at 28 °C. Four different treatments were tested with three replicates: (i) control (30 mL dH_2_O), (ii) bacteria treatment (30 mL *T. aidigensis* suspension, 10^8^ CFU/mL), (iii) Salt stress (30 mL 400 mM NaCl solution), and (iv) bacteria + salt combination (15 mL *T. aidigensis* (1×10^8^ CFU/mL) + 15 mL 400 mM NaCl]. Tomato seedlings were grown in the pots according to Çetinkaya et al. (2022) at 24°C for 21 days, and morphological development with plant growth involving root formation in all groups was visualised after rinsing the roots of the plants with distilled water at the end of the experiment (Lubna et al. 2025).

### Phytostimulation Assay /Pot Trials

Pot trials were carried out to determine the colonization ability of KB25 on the tomato rhizosphere (*Lycopersicum esculentum* Mill cv. Dünya). The tomato seedlings were grown in alluvial loam (total N 112; P 32.3; K 17.4; and Mg 9.1 mg/100 g soil; pH 6.5) at 20/15°C (16/8 h, day/night cycle). KB25 solution of 174 ml adjusted to 10^7^ CFU/ml was drenched at the 2-3 leaf stage (1 week) into tomato rhizosphere. The experiment was conducted with non-inoculated (control) and plants drenched with KB25 solution. Six weeks after planting of the tomato seedlings into the pots, 20 plants from each group with 5 replicates were randomly selected for measuring root growth and plant height. The experiments were carried out according to a completely randomized design and the data was statistically evaluated using the LSD test using SPSS (IBM SPSS Statistics for Windows, Version 22.0. Armonk, NY: IBM Corp.)

### Bioinformatics Analysis and Genomic Characterization

The bacterial samples were cultured in NB medium for 1 day, followed by centrifugation. DNA was purified from the pellet sections using the CTAB method (Nishiguchi et al. 2002). The purity of the extracted genomic DNA was assessed using 1% agarose gel. The purified DNA was used to construct a library with blunt triple junctions consisting of fragments of approximately 500 base pairs in length. Subsequently, the library was subjected to sequencing using an Illumina Hi-Seq 2500 sequencing system (BM Software Consultancy and Laboratory Systems Ltd., Türkiye).

We performed adapter trimming and quality filtering on the reads using Trim Galore version 0.6.7 (Krueger 2020), which incorporates Cutadapt version 3.5 (Martin 2011) and other parameters for cleaned read pairs was followed according to Bozkurt et al (2025). The resulting cleaned read pairs served as input for de novo assembly using SPAdes version 3.15.5 (Bankevich et al. 2012). The resulting scaffolds and contig were re-ordered against the reference genome of strain (NCBI acc. num; GCF_900215625.1). The final assembly was evaluated using QUAST 4.5. Annotation was performed using the PGAP pipeline and RAST (Rapid annotation using subsystem technology) (Brettin et al. 2015). The genomic sequence was also annotated using BLAST and compared with the UNiprot protein database (Bozkurt et al. 2025).

The total number of forward and reverse primer sequences were 3882623 (x2 for both strands) with Scaffold number 196. The final genome assembly comprised 134 contig with a total length of 3.882.623 bp. The longest segment spanned 140056 bp, and the N50 value was 44331N75 23115 with GC ratio: 41.6343. To identify the predicted genes, the assembled genome was subjected to Quast v. 5.0.2 analysis (Gurevich et al. 2013). Subsequently, PATRIC was used for further analysis (Wattam et al. 2017).

Protein sequences were annotated with Enzyme Commission (EC) numbers (Schomburg et al. 2004), Gene Ontology (GO) terms (Ashburner et al. 2000), and KEGG pathways (Kanehisa et al. 2016) using genome annotation pipelines (e.g., RAST and PROKKA) (Aziz et al. 2008; Seemann, 2014). ORF data obtained from the KB25 fna. The antibiotic production and bacteriocin genes were identified using the SEED viewer server (Aziz et al. 2012). An in-depth analysis of the antibacterial compounds produced by KB25 was conducted using AntiSMASH (version 6.0) to predict gene clusters involved in the production of secondary metabolites (Blin et al. 2021).

Of the resulting contigs, the 16S rRNA sequence was extracted and compared with reference genomes available in the databases. A phylogenetic tree was constructed using the neighbor-joining method with related sequences obtained from the BLAST output of NCBI. Gene profiles of the bacterial samples were created by examining the sequence data in the KEGG database, and species determination was performed by extracting 16S sequences from the sequence data. The entire genome of the bacterium (KB25) was mapped against the reference genome. A phylogenetic tree was constructed using the neighbor-joining method with related sequences obtained from the BLAST output of NCBI. Additionally, the KB25 gbk file was analysed for the average nucleotide identity (ANI) and DNA-DNA hybridisation (dDDH) using The Type (Strain) Genome Server (TYGS) (Meier-Kolthoff and Göker 2019). The fasta sequence of the whole-genome was submitted to PubMLST (https://pubmlst.org/), and allelic variation analyses were performed (Jolley et al. 2018).

### Dual Culture Experiment for Metabolite Extraction

To ensure metabolite production of *T. aidigensis* against *P. infestans*, three different testing groups were established: (i) KB25 strain was growth in NB culture at 28°C and 150 rpm for 18 hours alone. (ii) *P. infestans* was growth in PDB culture 28°C and 150 rpm for 24 hours alone. (iii) the supernatant of previous growth *P. infestans* in PDB culture was introduced into KB25s’ NB culture and incubated at 28°C and 150 rpm for 18 hours. Then, these three different cultures were centrifuged at 4000×g for 10 minutes and the resulting supernatants were used for further metabolite analysis (Liu et al. 2024). The bacteria were previously growth under salt stress conditions for all tested treatments before resulting supernatants.

### Metabolomics analysis

Biological samples of biofluids were collected and processed following standard protocols to ensure consistency and minimize degradation. Samples were stored at -20°C until analysis. Prior to mass spectrometry analysis, samples were thawed, homogenized, and processed using the appropriate extraction method for the targeted metabolites or proteins. In the case of metabolic profiling, metabolites were extracted using (Acetonitrile: Methanol: Milli-Q water; 40:40:20). Samples were subjected to vortex mixer for 2 min, sonication for 1 min and then centrifugation for 5 min at 14000×g and 4°C. The supernatants were collected in polypropylene tubes and dried in a nitrogen gas evaporator. The dried samples were suspended in 40 µL of extraction solvent (ACN: MeOH: MQ; 40:40:20) and vortexed for 2 min before being transferred into HPLC glass autosampler vials. 5 µL of samples were injected into a Bruker Elute LC system, coupled with a timsTOF pro 2 mass spectrometer (full scan, range: 20-1300 MS1: m/z and RT from inhouse standard library) that was equipped with heated electrospray ionization (H-ESI) source probe. The gradient elution was performed using 20 mM Formic acid (0.1%) acetonitrile (pH 2.7) with a flow rate of 0.25 mL/min. Formic acid solution (0.1%) was used as mobile phase A and Formic acid (0.1%) acetonitrile as mobile phase B. The gradient elution was done according to Online Resource 1. The instrument was controlled using Compass software (Bruker Elute LC system). The quality of the LC-MS runs was monitored by injecting pooled quality control (QC) samples made of EV samples and blank samples (extraction buffer) between the samples. The data was prefiltered for background (> 20% peak intensity in blank/sample) and response standard deviation (% coefficient of variation within pooled QC).

### Metabolomic data analysis

MZmine software (v4.7.3, Windows version) (http://mzmine.github.io/, downloaded on July 4, 2025) (Schmid et al. 2023) was utilized to preprocess the raw LC-MS spectra data. Firstly, the raw data was imported into the platform, and data processing was performed, including mass detection (centroid), chromatogram resolving, deisotoping, peak alignment, and filling in missing peak data. The noise level parameters were set for MS1 (2.0E3) and MS2 (3.0E2) before executing the ADAP chromatogram builder with a scan-to-scan accuracy set at 0.005 Da or 10 ppm, while keeping other parameters at default values. Finally, the resulting MS1 feature data was exported and save in Excel (.csv) file while the MS2 feature data was exported and saved in Topview version 3.4.1. (Sturm and Kohlbacher 2009).

The resulting peak intensity features of MZmine software was subjected to a statistical analysis including using the MetaboAnalyst 5.0 (Chong et al. 2019). The processed data was exported as CSV files containing information of the detected peaks, retention time and m/z values, to MetaboAnalyst 5.0. Data were normalized by sum, auto scaled and analysed using principal component analysis (PCA) and orthogonal partial least-squares discriminant analysis (OPLS-DA). For the PCA, five components were analysed to explain the variance between untreated and treated extracts. Then the OPLS-DA was used to further assess the significant difference between secondary metabolites extracted from the untreated and treated extracts. Multivariate data analyses tools were used to reveal signature metabolites that gratify the basis of p-value<0.05, false discovery rate (FDR) corrected p-values (q-values)<0.05, fold change (FC)>2.0, and variable importance in projection (VIP)>1.2. Cross-validation (CV) was performed using a tenfold CV method indicating the accuracy, Q2 and R2 values. A permutation test was also done to validate the model with the permutation p-value of p<0.01. Further functional characterization was performed by pathway enrichment analysis using MetaboAnalyst 5.0.

To confirm our findings on metabolomics data with genomic data of *P. infestans*, we have constructed integrated genomic-metabolomic response analysis using script R code (ver. 4.5.2) under Bioconductor (ver. 3.22). We used constructed matrix involving genome data and metabolomics data for 3 different treatments (*bacteria alone, bacteria+fungus*, and *fungus alone*) and the data were saved as.csv file and cointegrated using R code and converted into heatmap to show correlation between genomic results and metabolomic output.

### Molecular protein-protein and protein-ligand interactions

We detected the presence of the terpene cyclase SqhC gene from the whole genome sequence data of KB25. Using the amino acid sequence of this gene and its functional annotation data, we identified this gene sequence as sporulenol synthase (EC:4.2.1.137) via the KEGG orthology search database. Subsequently, based on our *in vitro* observations, we used sporulenol, a metabolite produced by the bacteria, as a ligand in molecular docking studies to investigate its interaction with both RxLR effector peptide protein of the pathogen and the glutamate receptor of plant. To see if the ABC-type proline/glycine betaine transport system, permease component, could replace the glutamate receptor of the plant when bacteria are present in the plant rhizosphere, an examination was conducted between the two proteins. The superposition of two aligned proteins is evaluated by the TM-score, calculated by the established method TM-align (Zhang and Skolnick 2005).

#### Modelling of ABC-type proline/glycine betaine transport system, permease component and glutamate receptor interaction

To investigate the interaction between ABC-type proline/glycine betaine transport system, permease component of strain KB25 (P_038566112.1 | COG1174) and GLR3.5 glutamate receptor of tomato (*Solanum lycopersicum* Gene ID: 101262276), we conducted computational analysis to validate the presence of unique sequences belonging to KB25 genomic annotation data. We retrieved the sequences of KB25 bacterial strain and translated to aminoacid sequences using ExPasy tool and resulting output was submitted to SwissMODEL for selecting the best matching model with high score (Gasteiger et al. 2003; Waterhouse et al. 2018). These data served as a base for modelling the corresponding ABC-type proline/glycine betaine transport system, permease component, which exhibited similarity with the active sites (A subunit) of ABC-type glycine betaine transport system substrate-binding domain-containing protein (A0A268HF21.1.A) and AlphaFold DB model of A0A268HF21_9BACI (gene: A0A268HF21_9BACI, (organism: *Terribacillus saccharophilus*) as templates for modelling SWISS-MODEL (http://swissmodel.expasy.org) was used considering Z-score.

In docking analysis, we selected the GLR3.5 glutamate receptor 3.5 protein of tomato (Gene ID: 101262276). Protein-protein docking was performed using the HADDOCK server (Table 1) (van Zundert et al. 2016). The active site/interface residues of ABC-type proline/glycine betaine transport system, permease component and GLR3.5 glutamate receptor 3.5 were given in Online Resource 1. The interacting residues were visualized using HADDOCK, PyMol and LigPlot+ (Schrödinger and DeLano 2023). The interaction was visualized using PyMOL Molecular Graphics System (ver. 3.1.3), integrated with Python version 3.12.4.

**Table 1.** Double-layer agar test results. **KB25 Treated**: Inoculated with *T. aidingensis* KB25. **Control**: The *P. infestans* group without bacterial inoculation. Asterisks indicate statistical significance (p<0.05)

|  |  |  | KB25 Treated |  |  | Control |  |  |
| --- | --- | --- | --- | --- | --- | --- | --- | --- |
|  |  |  | Mean | Std. Deviation | Std. Error Mean | Mean | Std. Deviation | Std. Error Mean |
| Statistic |  |  | 25.56* | 7.92 | 2.50 | 51.61* | 16.45 | 5.20 |
| Bootstrap<sup>a</sup> | 95% Confidence Interval | Lower | 25.56 | 7.92 |  | 51.61 | 16.45 |  |
|  |  | Upper | 25.56 | 7.92 |  | 51.61 | 16.45 |  |
| a. Unless otherwise noted, bootstrap results are based on 1000 stratified bootstrap samples<br>(* . The mean difference is significant at the 0.05 level according to L<sub>SD</sub>) |  |  |  |  |  |  |  |  |

#### Molecular Docking-based Virtual Screening for Protein-Ligand Interactions

The experimental X-ray diffraction structure of secreted RxLR effector peptide protein, putative (*Phytophthora infestans* T30-4), with its corresponding Protein Data Bank (PDB) identification number 7XP9, was obtained from the Research Collaboratory for Structural Bioinformatics (RCSB) PDB website (Berman et al. 2003; Webb and Sali 2016; Xing et al. 2023). The missing residues from the structure were added via PyMol’s builder plugin (open-source v2.5.0), and loop regions where the residues were added, which were refined using MODELLER (v10.1) (Schrödinger LLC 2023; Webb and Sali 2014). Subsequently, the structure was subjected to a purification process, in which all heteroatoms were removed, except for the atoms associated with cofactors. In addition, polar hydrogen atoms were introduced where deemed essential, and Kollman charges were calculated (van Zundert et al. 2016). A grid box measuring 16.6 Å in the X-axis, 16.6 Å in the Y-axis, and 16.6 Å in the Z-axis was computed to encapsulate the active site of the 7XP9 structure. Virtual screening was performed using the secreted RxLR effector peptide protein structural model, evaluating its interaction with the sporulenol ligand (PUBCHEMID: 71448999) within a pre-determined grid box. Screening was conducted with an exhaustiveness level of 64 using AutoDock Vina (v1.1.2) (Schrödinger and DeLano 2023). The ligands exhibiting the highest affinity scores were subsequently selected using the same configuration. The manually predicted bonds were also cross-validated using the TU Dresden’s Protein-Ligand Interaction Profiler (PLIP) webserver, and only overlapping interactions were considered (Schake et al. 2025). Furthermore, the similar docking procedures were carried out for A chain of the structure of glutamate receptor-like channel AtGLR3.4 (PDB: 7LZH) due to active binding site existence and with the same ligand (sporulenol).

## RESULTS

### Morphological plasticity in response to high salt stress and microscopic studies

The morphology of the microorganism was examined, and it was observed that increasing NaCl concentrations had a significant impact on cell structure (Fig. 1). Specifically, Gram staining and SEM images showed that as the salt concentration increased up to 10% NaCl, the cells gradually elongated. This morphological elongation suggests that the microorganism has developed an adaptive response to high osmotic pressure. The elongation of cells could provide physiological advantages, such as reducing membrane permeability or facilitating the maintenance of intracellular balance. Interestingly, at 10% NaCl, cells were observed to revert to a more rounded morphology.

**Fig. 1.**
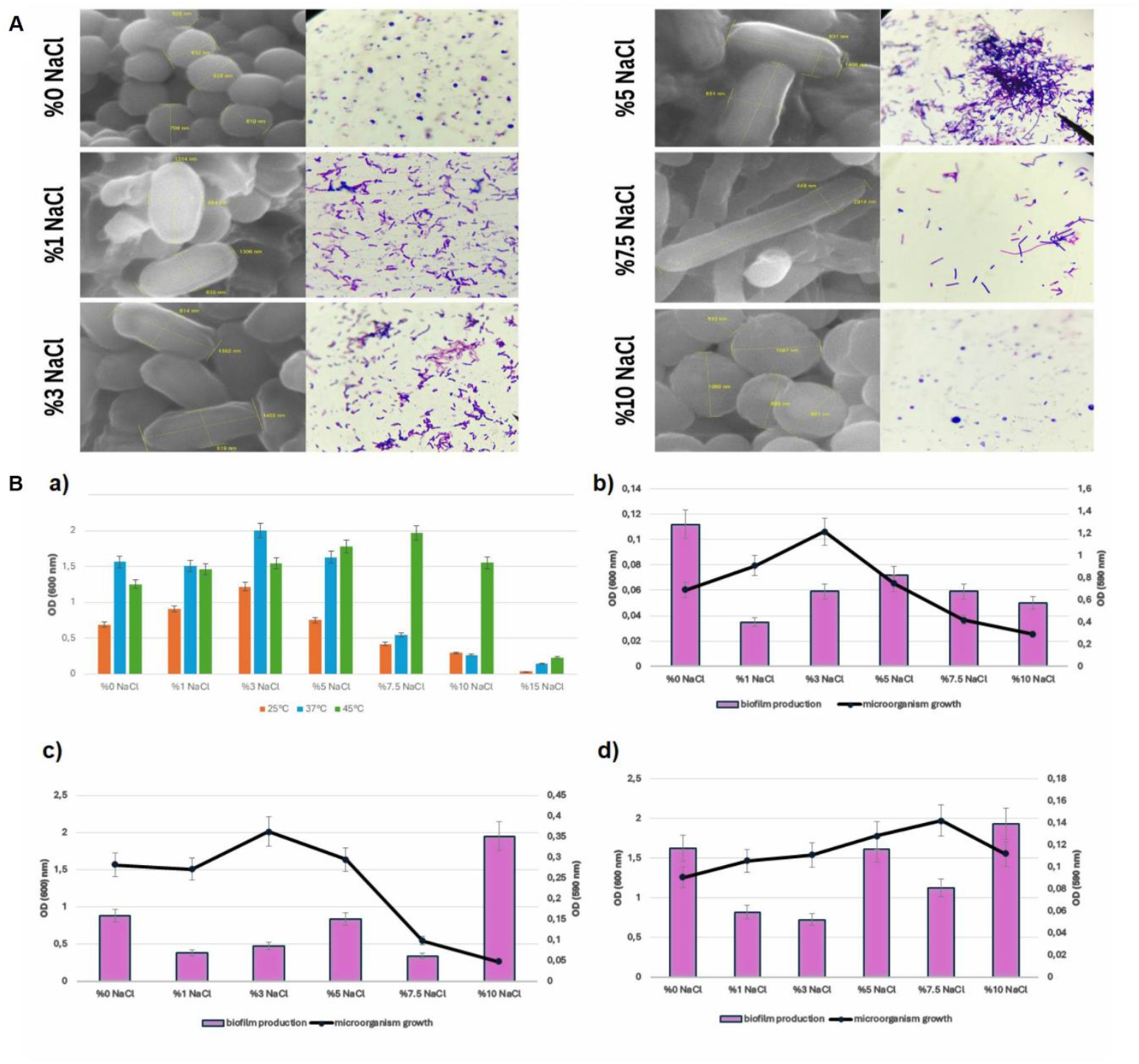
Morphological and physiological analysis of KB25 strain. (**A**) Morphological changes were visualized by SEM (magnification: 75kx), left) and Gram staining (right) at different NaCl concentrations. (**B**) **a)** Growth density of the halophilic microorganism under varying NaCl concentrations (0%, 1%, 3%, 5%, 7.5%, and 10%) at three different temperatures (25°C, 37°C, and 45°C). **b)** Effect of varying NaCl concentrations (0%, 1%, 3%, 5%, 7.5%, and 10%) on the biofilm formation potential of the halophilic microorganism at 25°C. **c)** Effect of varying NaCl concentrations (0%, 1%, 3%, 5%, 7.5%, and 10%) on the biofilm formation potential of the halophilic microorganism at 37°C. **d)** Effect of varying NaCl concentrations (0%, 1%, 3%, 5%, 7.5%, and 10%) on the biofilm formation potential of the halophilic microorganism at 45°C.

### The biochemical characterization of the KB25

The biochemical characterization of the KB25 isolate revealed positive reactions for catalase, urease, citrate utilization, lipase, putrefaction, and indole production. Additionally, the isolate was capable of fermenting both glucose and lactose with acid formation but without gas production, while the methyl red (MR) test yielded a negative result. This biochemical pattern suggests that the isolate possesses a metabolically versatile profile, capable of adapting to varying environmental conditions through both oxidative and fermentative pathways. Remarkably, the simultaneous presence of catalase, lipase, and indole production in a moderately halophilic bacterium has not been previously reported in the literature. This observation highlights the unique physiological and metabolic potential of the KB25 isolate, indicating that it may represent a novel or underexplored strain with valuable biotechnological properties. Therefore, these findings expand the current understanding of the biochemical diversity and adaptive mechanisms of halophilic bacteria, providing a new reference point for future enzymatic and genetic studies.

### The biofilm formation potential of KB25

The biofilm formation potential and growth capacity of a halophilic microorganism were comparatively evaluated under three different temperatures (25°C, 37°C, and 45°C) and six NaCl concentrations (0%, 1%, 3%, 5%, 7.5%, and 10%) (Fig. 1). The results demonstrated that both temperature and salt concentration significantly influenced biofilm production, with these effects often being closely associated with the microbial growth profile. At 25°C, the highest biofilm production was observed in the absence of salt (0% NaCl), whereas the highest growth was recorded at 3% NaCl.

### The Effect of Temperature on Antibiotic Susceptibility of the strain KB25

Temperature markedly influenced the antibiotic susceptibility profiles of moderately halophilic bacteria, revealing pronounced temperature-dependent variations (Online Resource 2). Among the antibiotics tested, imipenem (IPM), amoxicillin-clavulanic acid (AMC), and levofloxacin (LEV) were the most effective, showing inhibition zones of 48 ± 1.9 mm at 45°C, 44 ± 1.7 mm at 37°C, and 64 ± 2.4 mm at 25°C for IPM; 21 ± 1.1 mm, 17 ± 0.9 mm, and 40 ± 1.8 mm for AMC; and 33 ± 1.6 mm, 21 ± 1.2 mm, and 44 ± 2.1 mm for LEV, respectively. The superior efficacy at 25°C could reflect enhanced membrane permeability and antibiotic stability under milder thermal conditions, whereas partial activity retention at higher temperatures indicates robust β-lactam and fluoroquinolone performance. Conversely, antibiotics such as rifampicin (RA) and ofloxacin (OFX) exhibited their highest activity at 45°C (30 ± 1.3 mm and 31 ± 1.5 mm), which gradually decreased to 20 ± 1.0 mm and 25 ± 1.3 mm at 37°C and to 44 ± 2.0 mm and 38 ± 1.9 mm at 25°C, respectively, suggesting complex temperature-structure interactions influencing cell envelope permeability. Moxifloxacin (MOX) displayed also minimal activity only at 25°C (6.5 ± 0.4 mm) and was completely ineffective at elevated temperatures, implying thermal instability. Similarly, chloramphenicol (F) and cefepime (CEP) demonstrated their maximal inhibition at 25°C (30 ± 1.6 mm and 14 ± 1.0 mm) compared with reduced zones at 37°C (20 ± 1.2 mm and 25 ± 1.5 mm) and 45°C (26 ± 1.3 mm and 26 ± 1.6 mm, respectively). The efficacy of clindamycin (DA) and nitrofurantoin (NN) declined sharply with increasing temperature, from 24 ± 1.0 mm and 19 ± 0.9 mm at 25°C to 13 ± 0.7 mm and 0 mm at 37°C, and to complete inhibition loss (0 mm) at 45°C, indicating thermolabile antibiotic mechanisms. Moreover, gentamicin (CN) and amikacin (AK) maintained relatively consistent inhibition across the three temperatures (CN: 25 ± 1.2 mm, 22 ± 1.1 mm, 23 ± 1.0 mm; AK: 21 ± 1.4 mm, 15 ± 1.0 mm, 20 ± 1.2 mm), confirming the thermal resilience of aminoglycosides. Several antibiotics, including ceftriaxone (CRO), sulfamethoxazole-trimethoprim (CZ), azithromycin (AZ), and metronidazole (MTZ), exhibited negligible or no inhibition at any temperature (≤ 6 ± 0.5 mm), likely due to lack of specific targets or poor stability in saline environments. Meanwhile, netilmicin (NET) displayed a temperature-dependent response (20 ± 1.0 mm at 45°C, 9 ± 0.7 mm at 37°C, 10 ± 0.8 mm at 25°C), whereas piperacillin-tazobactam (TZP) showed an inverse pattern, increasing markedly from 29 ± 1.5 mm at 45°C to 14 ± 0.9 mm at 37°C and peaking at 45 ± 2.0 mm at 25°C, emphasizing the strong temperature adaptability and β-lactamase inhibition synergy.

### Antibiosis effect of strain KB25 using double-layer assay and pot experiments

In the double-layer assay, the antibiosis effect was significantly different depending on after inoculation period. Based on observation days, the suppressive effect of KB25 on the pathogen was found to be statistically significant from the 3 dpi to 7 dpi (Fig. 2, Table 1).

**Fig. 2.**
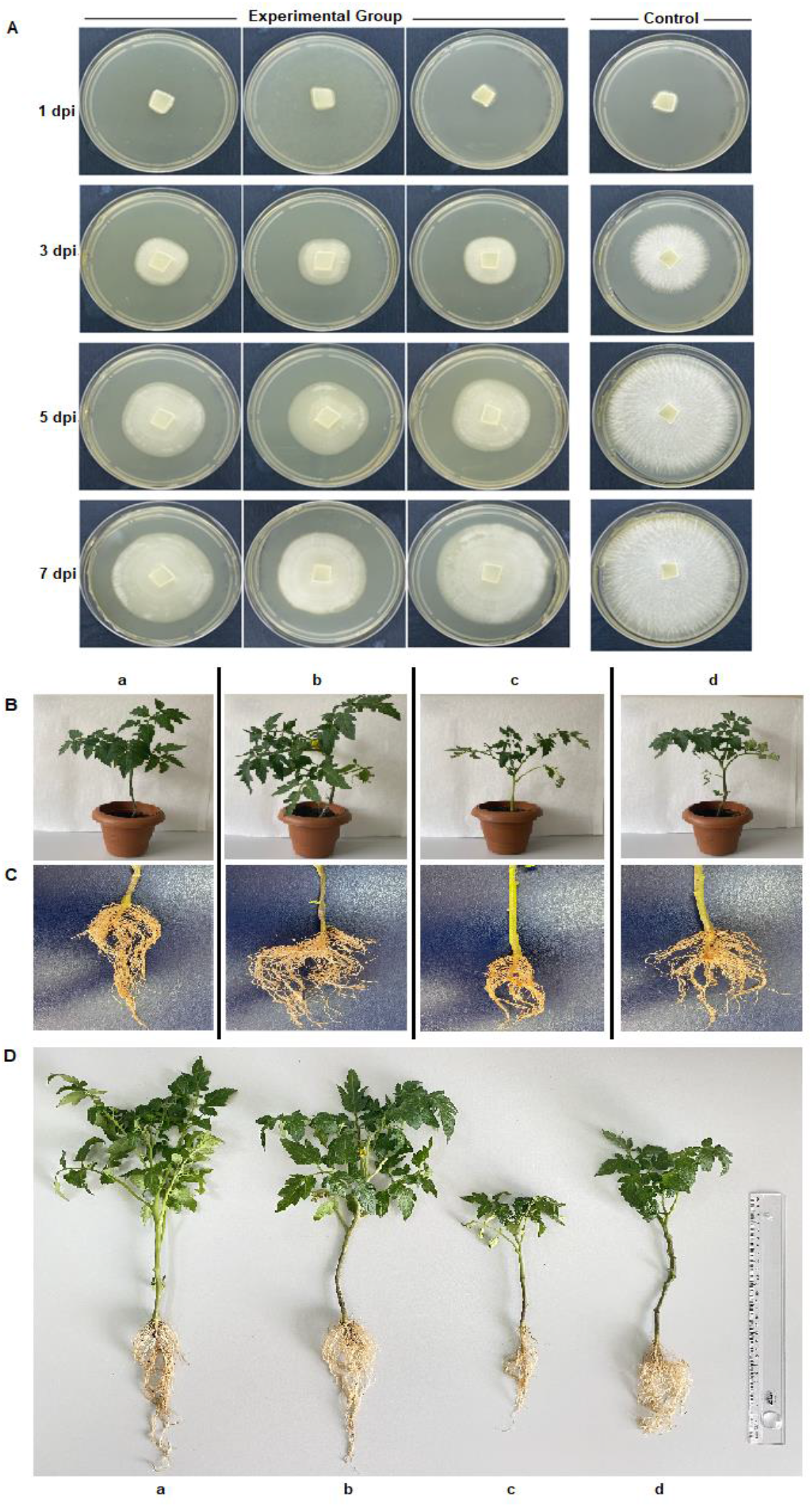
Pathogen inhibitory and salt stress regulation of strain KB25. (**A**) Double layer antagonistic assay to test inhibitory effect of KB25 against *P. infestans* according 7 dpi. (**B**) The pot experiment to test salt stress regulation effect of KB25 and, (**C**) root growth, (**D**) plant height after KB25 treatment compared to control groups under salt stress conditions **a**) Control, **b**) KB25 bacterial solution adjusted to 10^8^ CFU/mL drenched, **c**) Salt stress (400mM NaCl), **d**) Salt stress (400mM NaCl) + KB25 bacterial solution adjusted to 10^8^ CFU/mL drenched.

In pot experiments bacterial solution drenched groups have shown less salt stress which can be related with stress regulation effect of KB25. This effect was also confirmed with the root formation of bacteria treated plants that exhibited higher plant height compared to control groups and the plants exposed to salt stress alone during 21 days after treatment (Fig. 2).

### Genomic characterization of the bacterial strain KB25 showing superior suppression effect on *P. infestans*

The genome annotation resulted in the following taxonomy classification: cellular organisms Bacteria> *Bacillati*> *Bacillota*> *Bacilli*> *Bacillales*> *Bacillaceae*> *Terribacillus*> *Terribacillus aidingensis*. The Genomic size of KB25 is 3.882.623 bp, and comprises 4250 protein coding and GC content 41.6, (N50: 42.033, L50: 29, Number of contigs: 196, Number of subsystems: 270, Number of RNA: 103). Furthermore, we assigned 1938 sequences to ECs, 2849 sequences to GOGs, and 3540 sequences to KEGG pathways (Fig. 3A). The whole genome was annotated and compared with Pfam database (Online Resource 3).

**Fig. 3.**
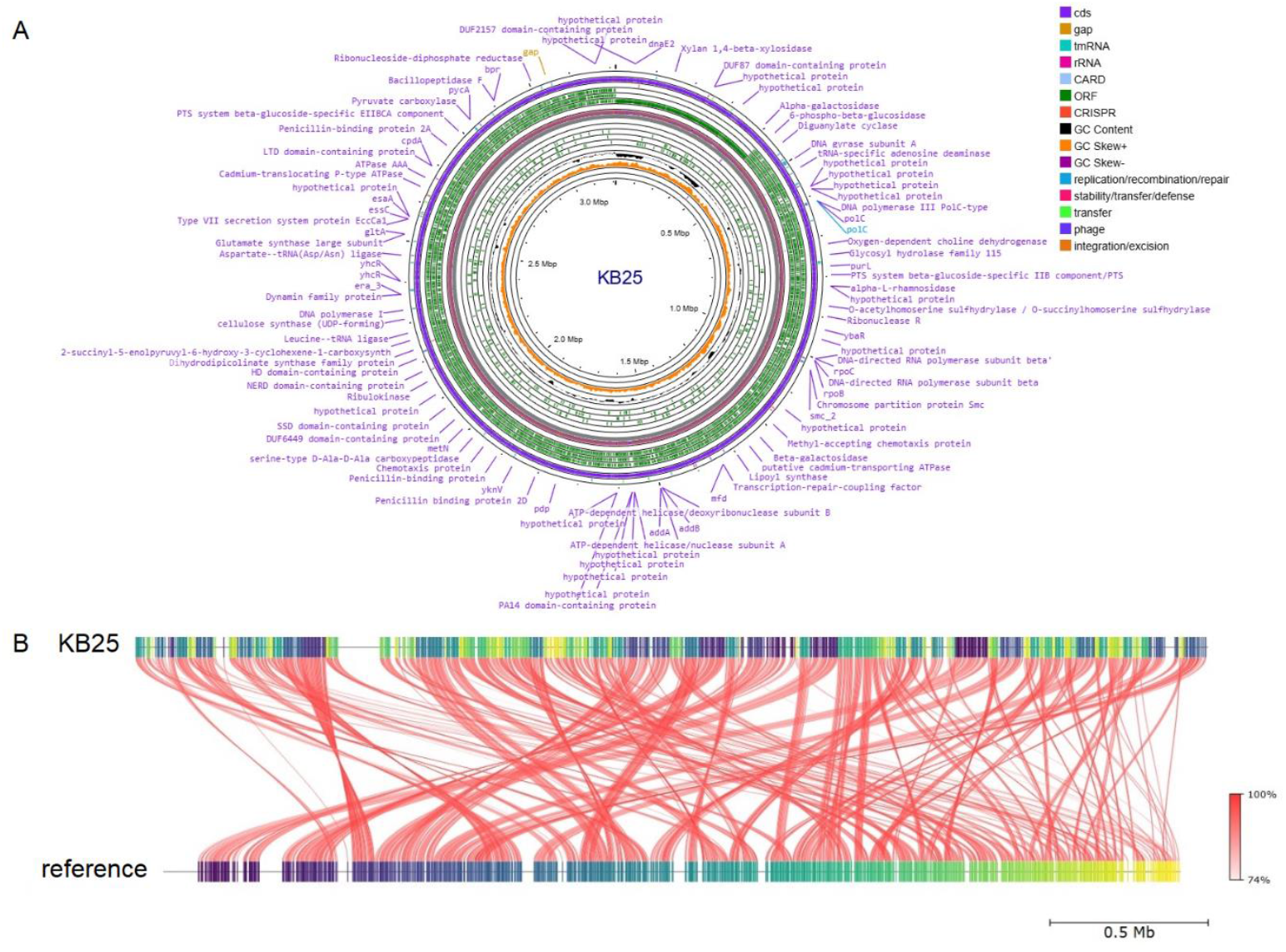
The figure shows genetical identification depending on whole genome sequence data of KB25. (**A**) The illustration shows a circular graphical display of the distribution of the genome annotations: CDS, tRNA genes, RNA genes, ORF regions, GC content, GC skew +/-, and predicted CARD genes in KB25 genome. (**B**) The data depicts the Average Nucleotide Identity (ANI) between KB25 and reference strain *T. aidigensis* CGMCC 1.8913.

Online Resource 4 presents the number of gene locations encoding proteins involved in specific biological functions and processes. The highest number of genes was identified for amino acids and derivates (212), whereas the lowest number was observed for Metabolism and aromatic compounds (3), potassium metabolism (3), cell division and cell cycle (3), and also, secondary metabolism (4), iron acquisition and metabolism (4) (Online Resource 4). No motility and chemotaxis genes were found which were correlated well with electron microscopy visualizations (Fig. 1). We observed the presence of genes encoding osmotic stress regulation; choline and betaine uptake and betaine biosynthesis (13), osmoregulation (1)(Online Resource 4).

16S rRNA sequences from genomic DNA were compared with reference sequences from NCBI for taxonomic analysis. Because of the comparison, KB25 (NCBI GenBank No: PX658357.1) showed the highest similarity with *T. aidigensis* CGMCC 1.8913 (NCBI acc. No. OBEK01000011.1, taxon no: 586416). In the phylogenetic tree prepared using the 16S rRNA gene, it was observed that the KB25 isolate was involved in the branching of *T. aidigensis* (Fig. 4C). Moreover, the Average Nucleotide Identity (ANI) (Fig. 3B), DNA-DNA hybridisation (dDDH)(Fig. 4A) and proteome analysis (Fig. 4B). Moreover, 16S rDNA results indicated that KB25 belongs to *Terribacillus aidigensis*. It seems genetically very close to the reference strain *T. aidigensis* CGMCC 1.8913. Neighbour-joining phylogenetic tree based on 16S rRNA gene sequences showing the relationship between *T. aidingensis* sp. nov. and members of the genus *Terribacillus* and other closely related genera. Bootstrap values (>50 %) based on 1000 replications are shown at branch nodes. Bar indicates 0.007 substitutions per site. The output and data were confirmed with de.NBI - German Network for Bioinformatics Infrastructure (Fig. 4C). rMLST analysis was also confirmed our previous results (Online Resource 5)

**Fig. 4.**
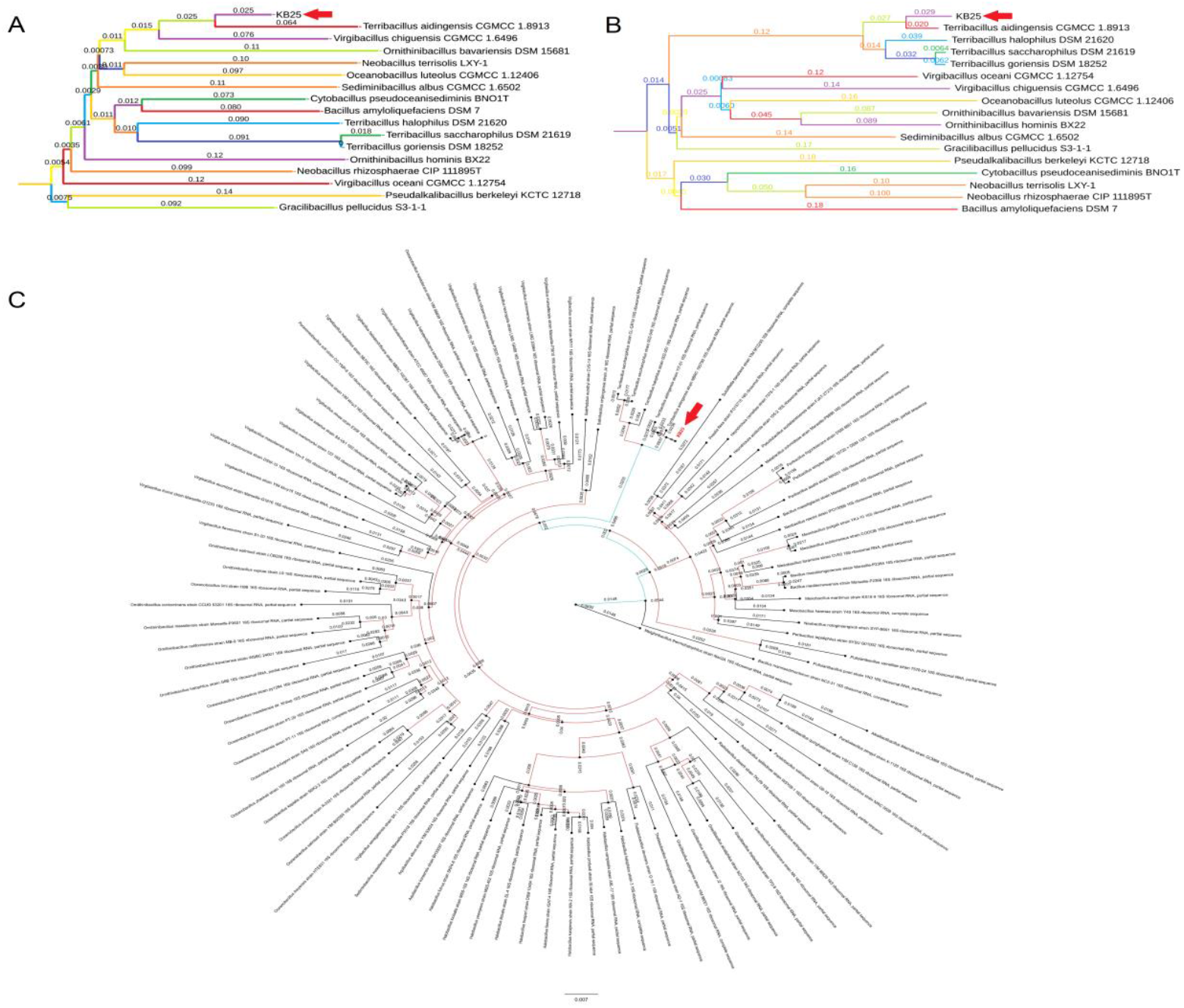
Phylogenetic analysis of strain KB25. (**A**) Genome-genome comparison. (**B**) Proteome comparison. (**C**) 16S rDNA analysis output using TYGS database. The red arrows indicated the place of our strain in various phylogenetic comparisons.

The whole genome sequence of KB25 was submitted to the antiSMASH database and compared with the other available gram-negative and gram-positive prokaryotes genomes deposited in this database. The results showed that the antimicrobial metabolites were only terpene derivates, as depicted in Online Resource 6. Furthermore, PathogenFinder results revealed that there are no pathogenic regions in our strain KB25.

### Genomic Functional Profiling and Protein Family Analysis of KB25

The genomic annotation of the bacterial strain KB25 identified 3,228 unique genes containing recognizable protein domains, with a total of 4.654 functional assignments. The profile is dominated by membrane transport systems, signal transduction machinery, and diverse metabolic enzymes, reflecting a genome adapted for environmental sensing and metabolic versatility. Dominant functional classes involve transport and signaling which are the most abundant protein family identified was the ABC transporter superfamily (n=92), alongside significant numbers of binding-protein-dependent transport systems (n=47) and major facilitator superfamily transporters (n=37). This extensive repertoire of transporters indicates a robust capacity for nutrient uptake (likely amino acids, sugars, and ions) and xenobiotic efflux, typical of organisms thriving in competitive or fluctuating environments. The analysis revealed a high prevalence of two-component system elements, including response regulators (n=33) and histidine kinase-like ATPases (n=30). The presence of these domains suggests the organism possesses intricate mechanisms to sense extracellular stimuli and rapidly modulate gene expression in response to environmental stress or nutrient availability. The metabolic machinery is characterized by a diversity of redox-active enzymes. Families such as short-chain dehydrogenases (n=28), aldehyde dehydrogenases (n=18), and aldo/keto reductases (n=13) are prominent. This abundance points to a flexible central carbon metabolism capable of managing diverse fermentation products and detoxification processes. Furthermore, acetyltransferases (n=30) were also highly represented. These enzymes are functionally diverse, playing roles ranging from peptidoglycan biosynthesis and antibiotic resistance to the regulation of cellular metabolism via protein acetylation.

The genomic profile of KB25 strain depicts a biologically versatile organism with a heavy investment in membrane transport and environmental sensing. The specific presence of quinone and lipid biosynthetic domains validates the metabolic activities observed in the metabolomics data, confirming a systemic link between the organism’s genetic potential and its functional metabolic output given below in metabolomics analysis part. The number of *T. aidingensis* KB25 genes related to molecular functions and cellular components, biological processes, comprehensive antibiotic resistance database outputs according to genomic data was given in Online Resource 7.

### Phytostimulation Assay /Pot Experiments

To directly test the plant growth effect of KB25 by *in vivo* conditions, we have growth the tomato plants in the soil inoculated with KB25 and non-inoculated plots. These *in planta* assays showed significant difference (p<0.05) in plant height and root formation of the plants inoculated with bacteria compared to untreated (control) ones Table 1. KB25 treatment gave rise to increased plant height and robust root growth under salt stress conditions (Fig. 2B, 2C, 2D).

### Metabolomics analysis

The visualizations demonstrated differences in the metabolome compositions in various samples including *bacteria alone* (Fig. 5A), *bacteria+fungus* (Fig. 5B) and *fungus alone* (Fig. 5C). A permutational multivariate analysis of variance (perMANOVA) showed the statistical significance of these differences (p<0.001). Changes in the abundances of individual metabolites indicate that most metabolites differ among *bacteria alone, bacteria+fungus, fungus alone* groups (Fig. 5). The metabolic diversity of various treatments supported by metabolome similarity was constructed using metorigin (ver. 2.0) software (https://metorigin.met-bioinformatics.cn/home/). A moderated t-test was performed to identify metabolites that exhibited a significant difference in abundance among *bacteria alone, bacteria+fungus, fungus alone* groups.

**Fig. 5.**
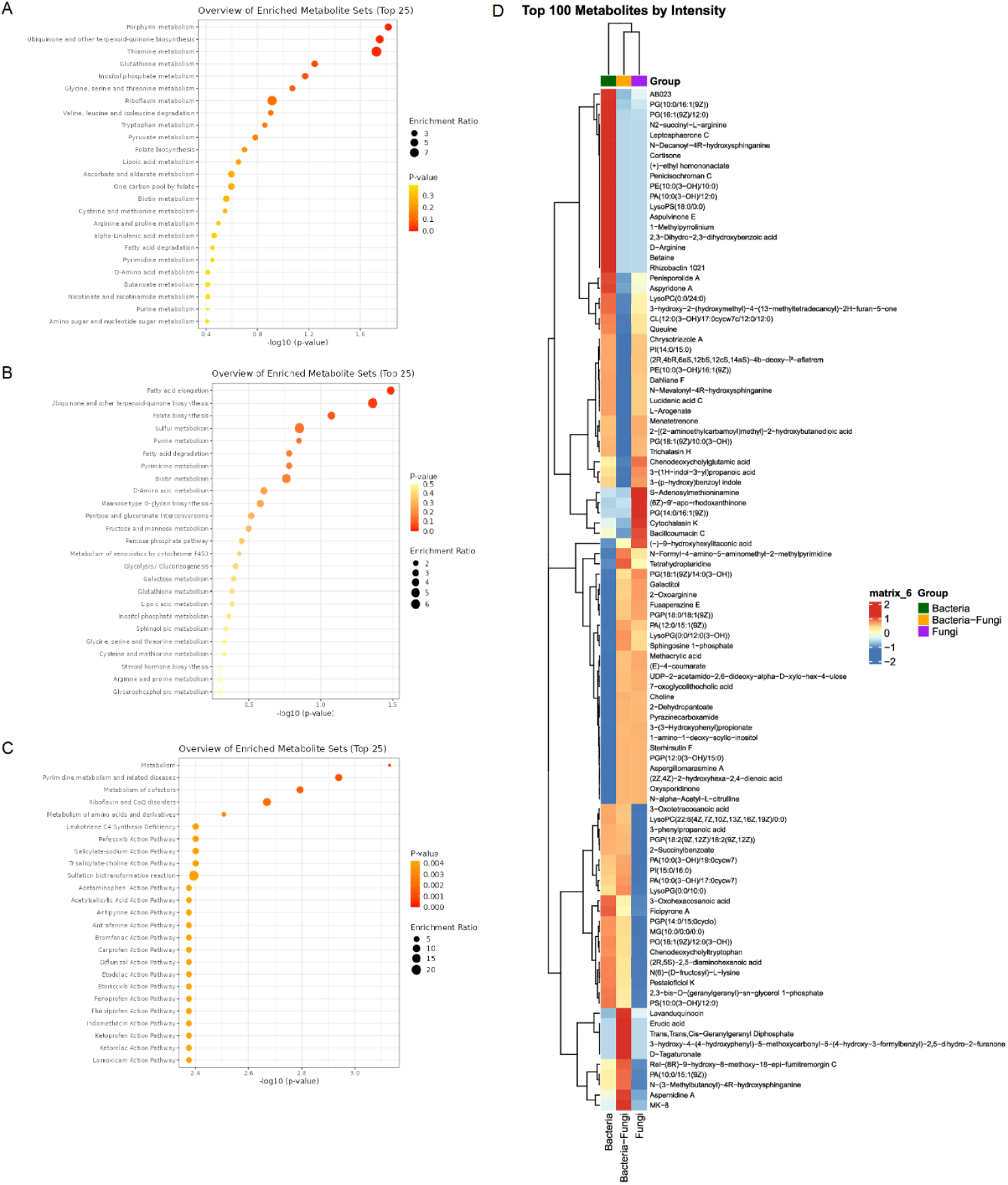
Metabolic Set Enrichment Analysis (MSEA) showing the overview of enriched metabolites sets in KB25 growth liquid culture. (**A**) *Bacteria alone*. (**B**) *bacteria+fungus*. (**C**) *fungus alone*. Metabolite sets enriched in various cultures were determined by over representation analysis (ORA) using the hypergeometric test (MetaboAnalyst 6.0). KEGG metabolic pathways containing at least 5 entries were used as a metabolite set library. The metabolite sets are ranked according to the p-value, and colour intensity from yellow to red indicates increasing statistical significance. The top 25 metabolite sets were demonstrated in the spot charts. The enrichment ratio represents the ratio of observed hits in the present data sets to expected hits in KEGG metabolic sets. (**D**) Heatmap of hierarchical clustering analysis of metabolite variations identified in the various liquid cultures within three groups, green colour represents *bacteria alone*, orange represents *bacteria+fungi* and purple represents *fungus alone*.

We performed metabolite set enrichment analysis (MSEA) using metabolites detected in our study and 25 top metabolic sets based on KEGG microbiome metabolic pathways as the metabolite set library to determine the enrichment of biologically relevant metabolic patterns in various treatments. The results from analysis are reported in Fig. 5 and Online Resource 8.

Three different extracts including *bacteria alone, bacteria+fungus, fungus alone* groups differed in terms of their metabolome. We visualised their top 100 metabolomic differentiation using a heat map according to abundance. We performed non-metric multidimensional scaling (nMDS) to ordain the samples based on a Euclidean distance matrix (Fig. 5D).

### Metabolic Pathway Enrichment Analysis

#### Bacteria alone

A metabolic pathway enrichment analysis was conducted to characterize the biological processes significantly associated with the observed metabolite profile. The analysis revealed profound alterations in cofactor metabolism specifically B vitamins and quinones alongside significant shifts in lipid and amino acid pathways. A total of nine pathways were identified as statistically significant (p*<*0.05). The most striking finding is the highly significant enrichment of vitamin-related pathways. Thiamine metabolism (p=1.08×10^-4^) emerged as the most significant pathway, driven by key intermediates such as thiamine monophosphate, thiamine triphosphate, and NAD. This suggests a critical upregulation of energy metabolism machinery, as thiamine pyrophosphate is a vital cofactor for enzymes like pyruvate dehydrogenase. Riboflavin metabolism (p=0.0051) was significantly enriched, evidenced by the presence of flavin mononucleotide. Furthermore, ubiquinone and other terpenoid-quinone biosynthesis (p=0.0033) showed significant activity, with metabolites like menatetrenone and phylloquinol identified. Collectively, the enrichment of thiamine, riboflavin, and quinone pathways points to a systemic enhancement of mitochondrial electron transport and oxidative phosphorylation capabilities. Glycerophospholipid metabolism (p=0.0018) was the second most significant pathway, characterized by a diverse array of lipids including choline, phosphatidylcholine, phosphatidylethanolamine, and phosphatidylserine. This indicates extensive remodeling of cellular membranes.

However, inositol phosphate metabolism (p=0.036) was also significant, involving signaling molecules like myo-inositol 1,4,5-trisphosphate and phosphatidylinositol-3,4,5-trisphosphate. This suggests that the observed lipid changes are not merely structural but likely involved in active signal transduction processes. The analysis also highlighted glycine, serine and threonine metabolism (p=0.0080), which bridges amino acid turnover with one-carbon metabolism via metabolites like 5,10-methylene-THF and choline. This is further supported by the significance of valine, leucine and isoleucine degradation (p=0.030), indicating active branched-chain amino acid catabolism for energy or biosynthetic precursors. Glutathione metabolism (p=0.017), represented by glutathione and oxidized glutathione, indicates an active cellular response to oxidative stress, aligning with the increased activity observed in the mitochondrial electron transport pathways.

Overall, the metabolomics profile of bacteria alone portrays a biological state characterized by intense metabolic activity, specifically driven by the upregulation of cofactor biosynthesis (thiamine, riboflavin, quinones) necessary for energy production. This is accompanied by significant membrane remodeling (glycerophospholipids) and active signaling (inositol phosphates), occurring against a backdrop of heightened oxidative stress management (glutathione).

#### Bacteria and Fungus Interaction

A total of eight pathways were found to be statistically significant (p<0.05). The most significantly enriched pathway was ubiquinone and other terpenoid-quinone biosynthesis (p=4.64×10^-4^), characterized by the presence of vitamin K_1_, phylloquinol, and alpha-Tocotrienol. This enrichment points towards a robust activation of electron transport chain components and antioxidant defense mechanisms, suggesting a heightened state of oxidative stress response or energy production. Closely related to energy metabolism is the significant enrichment of thiamine metabolism (p=0.0038) and pantothenate and CoA biosynthesis (p=0.036). Thiamine (vitamin B_1_) and coenzyme A are critical cofactors for the Krebs cycle and fatty acid metabolism. The identification of metabolites such as 5-aminoimidazole ribonucleotide and Dephospho-CoA suggests an upregulation of the machinery required for central carbon metabolism. Purine metabolism (p=0.0024) was highly significant, involving key intermediates like deoxyadenosine triphosphate and xanthosine. This indicates active DNA replication, repair, or signaling processes. Consequently, the one carbon pool by folate (p=0.0042) pathway was enriched, driven by folic acid and 5,10-methenyltetrahydrofolic acid. The synergy between folate and purine metabolism is well-established, as folate derivatives provide the necessary carbon units for purine ring synthesis, further supporting the hypothesis of increased cellular proliferation or turnover. While less dominant than the cofactor pathways, glycerophospholipid metabolism (p=0.019) showed significant perturbation, marked by choline and phosphatidylcholine. This likely reflects membrane remodeling events. Additionally, degradation pathways such as valine, leucine and isoleucine degradation (p=0.0095) and benzoate degradation (p=0.042) were enriched, with succinyl-CoA appearing as a common node. This convergence on succinyl-CoA reinforces the central role of the TCA cycle in assimilating carbon from diverse sources, amino acids and aromatic compounds to sustain cellular energy demands.

The metabolomics profile of bacteria and fungus interaction suggests a biological state characterized by upregulated energy production and cofactor biosynthesis (quinone, thiamine, CoA), coupled with active nucleotide synthesis (purine, folate). The data collectively point towards a system undergoing rapid growth, active repair, or a significant metabolic shift requiring enhanced mitochondrial function and carbon flux through the TCA cycle.

#### Fungus Alone

The analysis on sample obtained from fungus alone treatment revealed distinct alterations in lipid metabolism, energy-related biosynthesis, and secondary metabolite pathways. A total of seven pathways were identified as statistically significant (p<0.05). The most substantial metabolic shift was observed in glycerophospholipid metabolism (p=1.09×10^−4^), which showed the highest degree of enrichment. This pathway was characterized by the significant perturbation of key lipid species, including choline, phosphatidylcholine, phosphatidylethanolamine, and phosphatidic acid. The prominence of this pathway suggests a potential remodeling of cellular membranes or alterations in lipid signaling cascades. Arachidonic acid metabolism (p=0.027) was also significantly enriched, driven by metabolites such as leukotriene B4 and phosphatidic acid. Together with the findings in glycerophospholipid metabolism, this points towards a broader inflammatory or signaling response involving lipid mediators. Fatty acid elongation (p=0.0053) was also identified as a significant process, further supporting the central role of lipid dysregulation in the studied condition. Beyond lipid metabolism, the analysis highlighted significant activity in ubiquinone and other terpenoid-quinone biosynthesis (p=2.29×10^−3^). This pathway, which involves metabolites like menatetrenone and all-trans-nonaprenyl diphosphate, is critical for electron transport and cellular respiration, indicating potential impacts on mitochondrial function or oxidative stress responses. Purine metabolism (p=0.047) and phenylalanine, tyrosine and tryptophan biosynthesis (p=0.041) were also found to be significant, suggesting localized disruptions in nucleotide turnover and amino acid biosynthesis, respectively. The flavone and flavonol biosynthesis pathway (p=0.0055) were significantly enriched, driven by isoquercitrin and astragalin. Depending on the microbiome metabolism, this may indicate active secondary metabolite synthesis or the metabolic processing of dietary polyphenols.

The metabolomics profile of fungus alone is dominated by significant perturbations in lipid-related pathways, specifically glycerophospholipid and arachidonic acid metabolism alongside notable shifts in quinone biosynthesis. These results indicate a physiological state characterized by membrane lipid remodeling and potential alterations in energy metabolism.

The analysis mathematically correlated that the KB25 strain produces antifungal terpene derivatives under salt stress. The fact that the sporulenol synthase gene and the sporulenol/terpene peaks appearing on the heatmap in LC-MS/MS are in the same group (*bacteria+fungus*) and have a dark red (positive correlation) relationship with each other. This indicates the metabolite is not produced by spontaneously, but by the genes belonging to KB25. Our study also showed that the ABC-type permeases, we previously mentioned, are not only present but also stimulated simultaneously with the transport of osmolytes such as betaine and proline when encountering with the fungus. These findings suggest an integrated mechanism at which the bacteria both protects itself and attacks *P. infestans*. With this analysis, we have scientifically confirmed that KB25’s antifungal and halotolerance (salt resistance) capabilities have moved beyond being mere potential abilities and have become actual biodefense agent in the presence of fungus.

#### Molecular protein-protein and protein-ligand interactions

As known in bacteria the periplasmic binding protein acts like a clam shell that snaps shut around a nutrient (like proline or glycine betaine) and delivers it to the permease (the channel) to pump it into the cell. In plants, the plant glutamate receptor has this same clam shell domain floating outside the membrane. When it catches an amino acid (like glutamate), it pulls the channel open to let calcium (Ca^2+^) into the cell.

Based on our *in vitro* experimental results, we hypothesized that KB25’s ability to regulate salt stress is dependent on a number of metabolites it produces. To understand the potential for the bacterium’s metabolites to be used in the plant, we examined the interaction between two proteins responsible for stress regulation in terms of structural homology. Furthermore, when we investigated the possibility of these two proteins whether they substitute within each other using both TM-align and FoldMASON-MS-alignment tools, we found matching regions (Online Resource 9). On the other hand, as known, both the bacterial ABC transporter and the plant glutamate receptor proteins share the clam shell mechanism in their protein structure (O’Hara et al. 1993). Given this profound structural homology, it is hypothesized that the bacterial protein may exhibit functional mimicry or complementation of the plant receptor (Chiu et al. 1999). Such molecular crosstalk would be advantageous for rhizospheric bacteria, allowing them to modulate host signalling pathways to regulate plant stress responses.

While the bacterial system utilizes this domain for the uptake of osmoprotectants (proline/glycine betaine) to confer direct osmotic resistance, the plant GLR has adapted this domain to sense environmental amino acids, translating metabolic cues into cytosolic Ca^2+^ signalling cascades that regulate systemic drought and salt stress responses (Yu et al. 2022; Grenzi et al. 2022). The metabolites produced by KB25 could be involved in the stimulation of the glutamate receptor as reported by Price et al (2012). In this case, it increases Ca^2+^ influx, and in our *in vitro* pot experiments, we observed that stem thickening, and possible protection against osmotic pressure with root elongation and development in plants indicated that KB25 regulates salt stress.

The docking simulation reveals a stable heterodimeric complex characterized by a substantial interface area between ABC-type proline/glycine betaine transport system, permease component of strain KB25 (blue) and glutamate receptor (red). The surface representation (Fig. 6A) illustrates a high degree of shape complementarity, suggesting a tight steric fit between the two subunits. The ribbon diagram (Fig. 6B) further elucidates the secondary structural elements involved in binding, indicating that the interface is formed primarily through loop regions and surface helices, which facilitate the orientation of the two globular domains. Detailed inspection of the interface (Fig. 6C) reveals a dense network of non-covalent interactions. The contact surface is characterized by the close packing of side chains, minimizing solvent accessibility at the binding core. To deconstruct the specific atomic contributions to binding affinity, we analysed the 2D interaction map generated via LigPlot+ (Fig. 6D). We identified approximately 11 distinct hydrogen bonding pairs bridging the interface. The data indicated that Arg207(A) serves as a pivotal anchor, forming hydrogen bonds with both Asp51(B) and Lys298(B). Another a strong electrostatic interaction is observed between Arg239(A) and Glu284(B). At the N-terminal region of ABC-type proline/glycine betaine transport system, permease, Asp4(A) and Glu2(A) interact with Gln75(B), stabilizing the peripheral region of the interface. Tyr231(A) acts as a hydrogen bond donor/acceptor with Glu50(B), further securing the central contact region. The binding affinity was -13.9 kcal/mol (Online Resource 10).

**Figure. 6.**
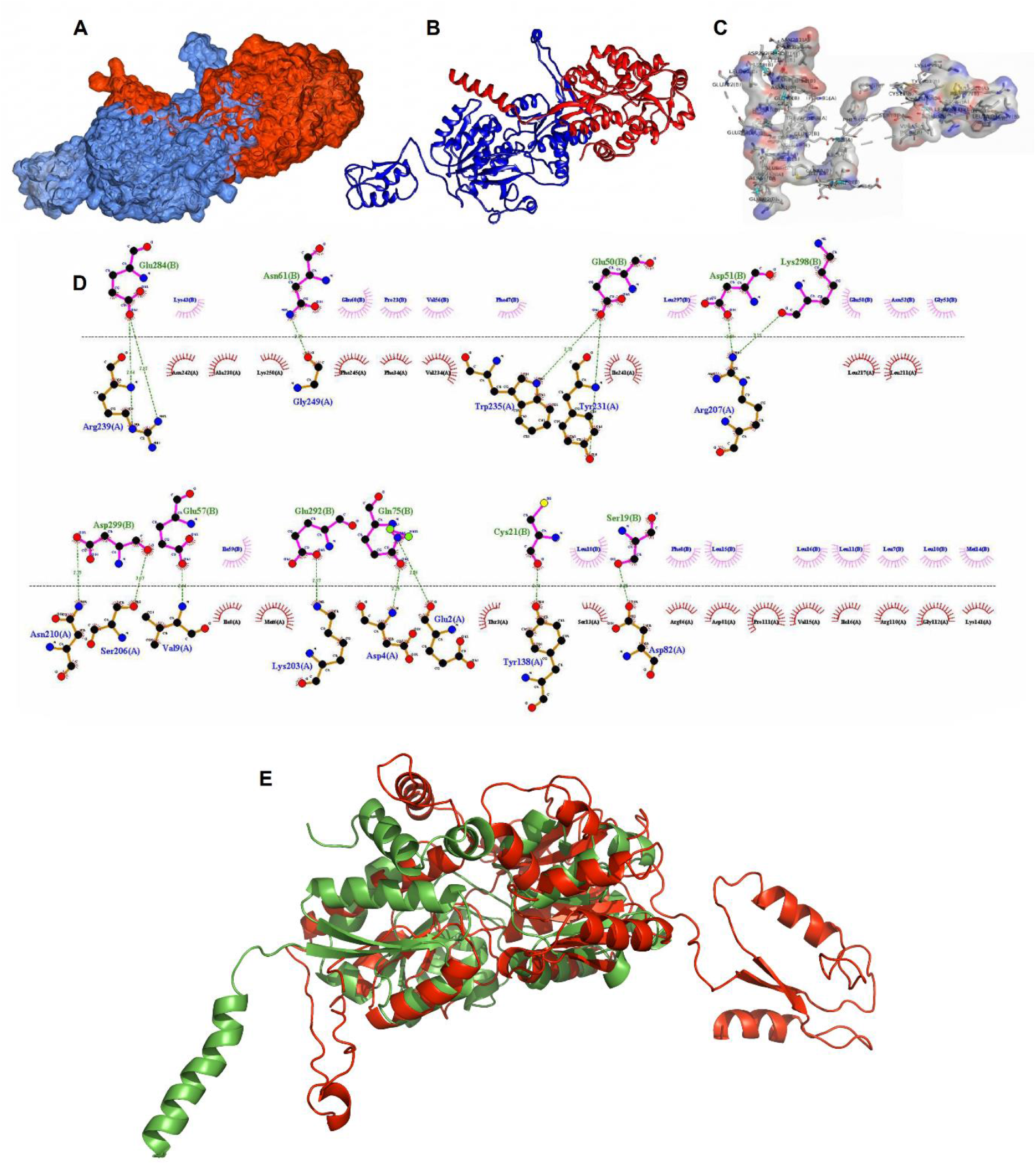
Molecular protein-protein docking results of ABC-type proline/glycine betaine transport system, permease component of strain KB25 (blue) and glutamate receptor (red). (**A**) Surface covered of protein-protein interaction. (**B**) Protein-protein interaction considering molecular surfaces. (**C**) Protein-protein interaction with hot spots colored red against the otherwise grey surface. (**D**) Protein-protein interaction analysis results computed by Ligplot+. (**E**) TM-align results of ABC-type proline/glycine betaine transport system, permease component of strain KB25 (green) and glutamate receptor (red).

The interaction map highlights a distinct hydrophobic patch, particularly rich in leucine residues on glutamate receptor. Leu7, Leu10, Leu11, Leu15, Leu16, and Leu18 on Chain B form a hydrophobic cluster that interdigitates with non-polar residues on ABC-type proline/glycine betaine transport system, permease, including Val15, Ile16, and Pro111. The structural model demonstrates a high-affinity interaction driven by a hydrophobic core (specifically the Leucine-rich motif on glutamate receptor) and stabilized by peripheral salt bridges and hydrogen bonds (specifically involving Arg207 and Arg239 on ABC-type proline/glycine betaine transport system, permease).

These six figures characterize the structural basis for the binding of the polycyclic ligand sporulenol to its target receptors. In Fig. 7B the most notable specific interaction is a strong hydrogen bond between the ligand’s oxygen atom and Lys96(A). The bond length is 2.51 Å, which is very short and indicative of a highly stable, possibly charge-assisted hydrogen bond (given Lysine’s positive charge). This single point likely orients the entire molecule. The rest of the binding energy is derived from extensive van der Waals contacts (indicated by the red eyelashes). The ligand interacts with a suite of aliphatic and aromatic residues: Ile10, Leu14, Phe100, Ile142, and the aliphatic portions of Arg101 and Arg146. Residue Lys93(A) is explicitly drawn (bottom orange bonds) rather than just shown as a contact, suggesting it forms a significant close-contact interface with the alkyl tail of the sporulenol, essentially capping the binding site.

**Figure. 7.**
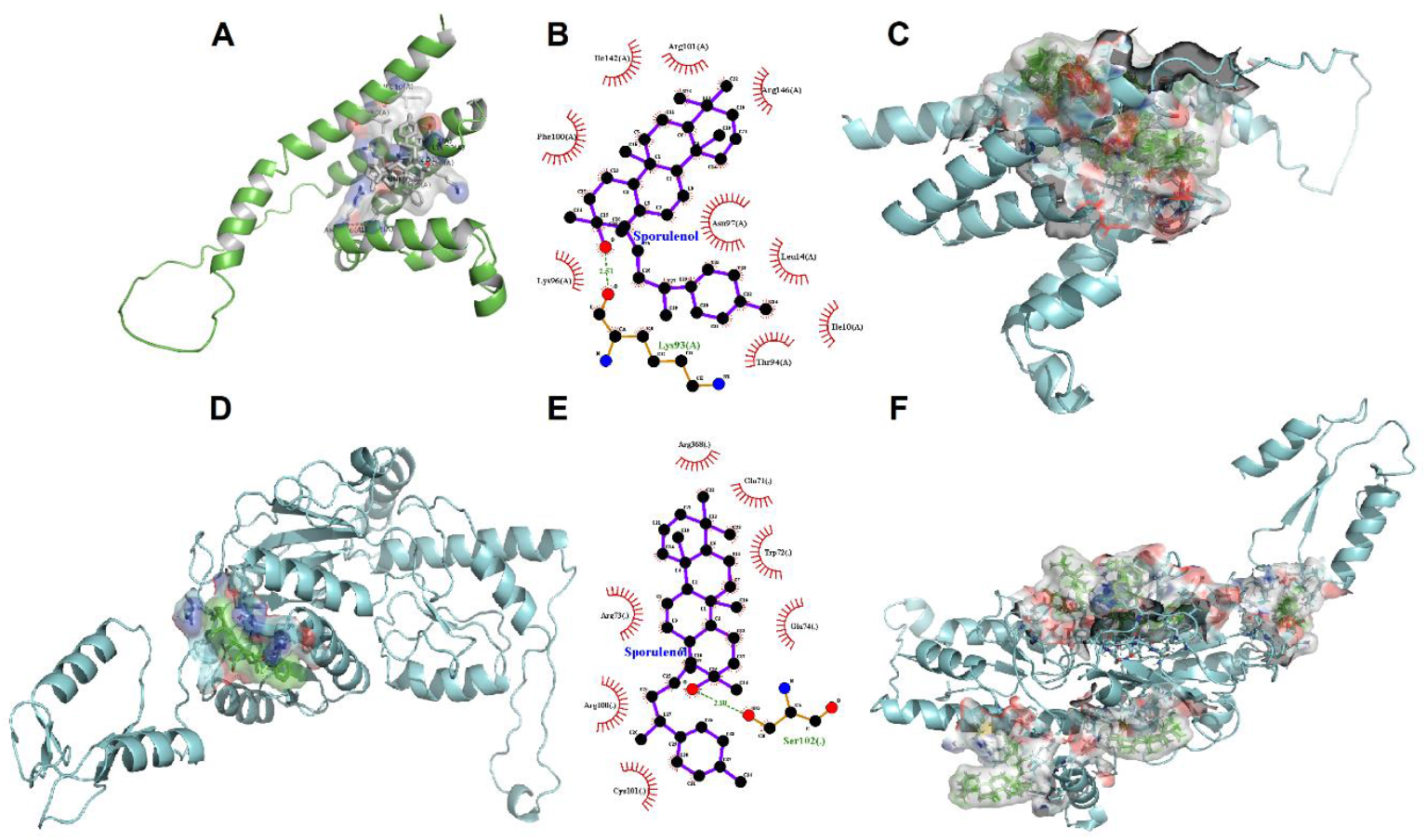
Molecular protein-ligand docking results of two target proteins, RxLR effector peptide protein, putative (*P. infestans* T30-4)(**A-B-C**) and glutamate receptor (**D-E-F**) with sporulenol as the ligand. (**A-D**) Protein-ligand interaction with hot spots colored red against the otherwise grey surface. (**B-E**) Protein-ligand interaction analysis results computed by Ligplot+. (**C-F**) Protein-ligand interaction considering molecular surfaces. (**F**) The complete docking conformation of target proteins, focusing multiple sporulenol compound binding events.

**Fig. 8.**
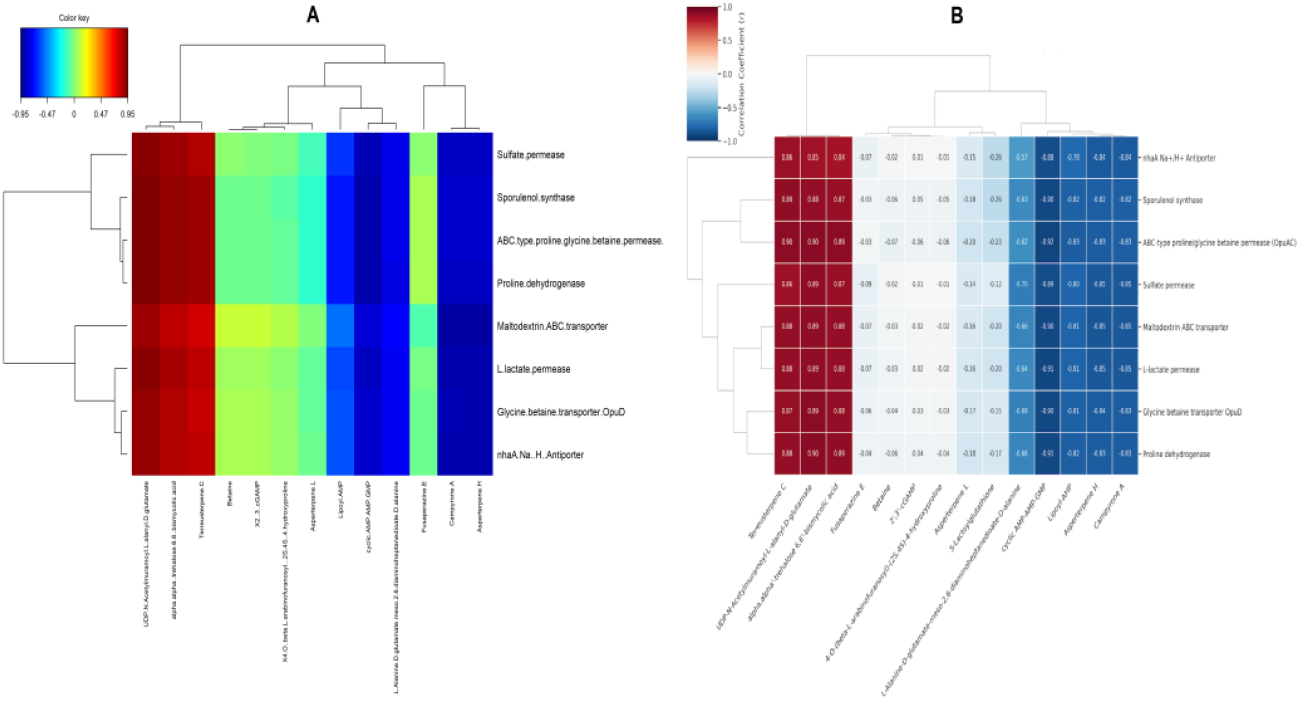
Integrated genomic-metabolomic response of KB25. (**A**) Heatmap depicts the top correlated genes and metabolomics of KB25. The correlation increases between genome and metabolomics data from dark blue to dark red (-0.95 to 0.95). (**B**) Clustered image map related to gene-metabolite integration depending on correlation coefficient values of compared genes and metabolites. The correlation increases between genome and metabolomics data from dark blue to dark red. (-1 to 1). For both heatmaps right side (y-axis) indicates the top metabolites with correlated genes in KB25 genome (x-axis).

Generally, the docking simulation reveals that sporulenol binds to a deep, hydrophobic cleft in the receptor. The complex is stabilized by a lock-and-key mechanism where Lys96 serves as the electrostatic lock (2.51 Å H-bond) and the surrounding hydrophobic residues (Ile, Leu, Phe) provide the shape-specific keyhole. The highest binding energy between RxLR effector peptide protein, putative (*P. infestans* T30-4) and ligand sporulenol was -7.06 kcal/j (Online Resource 11).

The Fig. 7D shows the tertiary structure of the target protein glutamate receptor (cyan ribbon) with the ligand sporulenol docked in a surface-accessible cavity. In Fig. 7E, the interaction was between specific hydrogen bond and the ligand coordinated with Ser102. The bond length is 2.80 Å, a canonical distance for a stable hydrogen bond, serving as the primary directional constraint for the ligand. This binding site is rich in charged and polar residues, specifically Arg73, Arg100, Arg368, Glu71, and Glu74. These results define a surface-accessible binding site for sporulenol anchored by Ser102. The highest binding energy between glutamate receptor and ligand sporulenol was -6.55 kcal/j (Online Resource 12).

## DISCUSSION

*In vivo* experiments demonstrated that *T. aidigensis* strain KB25 confers enhanced salt tolerance to tomato plants. Concurrently, microscopic analysis revealed distinct morphological adaptations of the strain when subjected to saline environments. Based on these observations, we suggested that strain KB25 exerts an inhibitory influence on *P. infestans* within plants challenging salinity stress. To validate this, the antagonistic potential of KB25 against *P. infestans* was assessed via dual culture assays, which confirmed significant suppression of the pathogen. Subsequent genomic characterization identified specific gene clusters within KB25 associated with stress regulation, as well as biosynthetic pathways for sporulenol, a terpene derivative. To elucidate the chemical basis of the observed *in vitro* suppression, cell-free culture filtrates from salt-stressed KB25 were introduced to *P. infestans* liquid cultures. Comparative metabolomic profiling conducted throughout the pathogen’s growth phase conclusively attributed the inhibitory activity to sporulenol. Further molecular investigations focused on protein-protein interactions to delineate the interplay between the KB25 derived ABC-type proline/glycine betaine transport permease implicated in osmotic adjustment and the plant glutamate receptor involved in salt stress modulation. We attempted to investigate the compatibility between osmotic gene products produced by bacteria and proteins that regulate osmotic stress in plants, examining the probability of one replacing the other using protein-to-protein similarity analysis. Finally, molecular docking simulations were employed to model the structural interactions of sporulenol with two key targets involving the pathogen’s virulence-associated RxRL effector protein and the plant stress-regulatory glutamate receptor.

The morphological plasticity observed in this study highlights a critical adaptive strategy of our strain KB25 to osmotic stress. The elongation of cells in response to increasing NaCl concentrations (up to 10%) aligns with recent findings on halophilic stress responses, where filamentation serves to increase the surface-to-volume ratio, thereby enhancing nutrient uptake and distributing osmotic pressure across a larger cell envelope (Saini et al. 2023). The biochemical profile of the KB25 isolate demonstrates a rare metabolic versatility. While halophilic bacteria often specialize in either oxidative or fermentative metabolism, KB25’s ability to ferment glucose and lactose (acid positive, gas negative) while maintaining strong oxidative traits (catalase, citrate) indicates a facultative anaerobic capability essential for fluctuating saline environments.

The uncoupling of bacterial growth and biofilm formation at 25°C where biofilm peaked without NaCl despite optimal bacterial growth occurring at 3% NaCl supports the stress-induced biofilm hypothesis. This finding suggests that hypoosmotic stress triggers biofilm pathway activation as a protective response (Ning et al. 2023). Furthermore, temperature exerted a profound modulatory effect on antibiotic efficacy (Richardson 2023; Van Eldijk et al. 2024).

The observed morphological changes in the microorganism under varying NaCl concentrations provide compelling evidence of its adaptive strategies to osmotic stress. As demonstrated by Gram staining and SEM analyses (Fig. 1), cells exhibited progressive elongation with increasing salt levels up to 10% NaCl. This elongation aligns with known halotolerant responses in various bacterial species, where cell shape modifications serve to mitigate the effects of hyperosmotic environments (Vauclare et al. 2020; Gandhi and Shah 2016). For instance, similar elongation has been reported in halophilic archaea and certain Gram-positive bacteria, such as *Bacillus subtilis*, under saline stress, potentially enhancing surface-to-volume ratios to optimize ion transport and osmolyte accumulation.

This property could facilitate the intracellular accumulation of compatible solutes, such as glycine betaine to counterbalance osmotic pressure and maintain turgor. Such mechanisms are consistent with broader microbial physiology, where morphological plasticity enables survival in fluctuating salinity habitats (Yu et al. 2024). These findings underscore the microorganism’s potential as a model for studying osmoregulation and have implications for biotechnological applications, including salt-tolerant probiotics or bioremediation in hypersaline wastewaters.

The biofilm formation and growth capacity of the halophilic microorganism were significantly influenced by temperature and NaCl concentration (Fig. 1). For instance, in *Halomonas meridiana, Kushneria indalinina*, and *Halomonas aquamarina*, maximum biofilm formation occurred at various NaCl concentration up to 1 M (Qurashi and Sabri 2012). The similar results were observed on *Tetragenococcus halophilus* near its optimal growth range (5-10% NaCl), with temperature strongly modulating capacity (Yao et al. 2022). In contrast, our results at 25°C favor biofilm at zero added salt, possibly reflecting an energy-efficient adaptation to combined mild osmotic and thermal stress. These findings collectively highlight that temperature exerts a dual influence on antibiotic efficacy directly by altering drug stability and indirectly by modulating bacterial physiology. Collectively, the data emphasize that the antibiotic susceptibility of moderately halophilic bacteria is influenced not only by the molecular nature of the antimicrobial agent but also by environmental temperature. Therefore, antimicrobial susceptibility testing in extremophilic microorganisms should consider non-standard conditions, including environmental stress-factors such as temperature.

The application of taxonomic approach, integrating 16S rRNA phylogeny with whole-genome metrics (ANI and dDDH), has definitively classified isolate KB25 as *Terribacillus aidingensis*. This genomic validation is critical, as recent studies emphasize that phenotype-based classification alone is often insufficient for distinguishing closely related halophilic species (Chun et al. 2018). In our investigation, we realized that there is still no complete clarity regarding the taxonomy of *Terribacillus* species. We identified our strain as *T. aidingensis* by examining our annotated data according to genome-to-genome comparisons. Furthermore, the 16sRNA result and RMLST result confirmed our identification (Jolley et al. 2012).

The significant alleviation of salt stress observed in the pot experiments provides functional validation for the genomic presence of choline and betaine uptake systems. The accumulation of compatible solutes like glycine betaine is a primary mechanism by which halotolerant bacteria maintain cellular turgor under hyperosmotic conditions without perturbing central metabolism (Nadeem et al. 2013). KB25 likely confers induced systemic tolerance to the host plant, a phenomenon where bacterial modulation of the root environment enhances the plant’s own stress response machinery which is accordance with our pot experiments. The improved root formation and plant height in KB25-treated groups further indicate that this protection extends plant survival under salt stress conditions.

While the double-layer assay demonstrated significant antibiosis (3-7 days post-inoculation), genome mining via antiSMASH surprisingly identified terpene derivatives as the sole antimicrobial class. While less characterized than polyketides or lipopeptides in bacteria (Caulier et al. 2019), recent genomic surveys of aerobic endospore-forming bacteria have highlighted the ecological role of volatile terpenes in competitive exclusion and defense against fungal pathogens (Yamada et al. 2015; Weisskopf et al. 2021). Furthermore, the PathogenFinder results confirming the absence of pathogenic genomic regions establish KB25 as a biosafe candidate for agricultural application.

The significant promotion of tomato plant height and root proliferation under salt stress provides *in vivo* validation of the plant growth-promoting traits identified in our genomic and *in vitro* assays. The robust root growth likely results from the bacterial modulation of auxin/cytokinin ratios or the direct provision of compatible solutes, which lowers the osmotic potential of root cells to facilitate water uptake as described in previous studies (Nguyen et al. 2025; Liu et al. 2022).

The metabolomic profiles reveals that the interaction between KB25 and the fungal pathogen triggers a specific metabolic formation. In the *Bacteria Alone* group, the enrichment of vitamin B pathways specifically thiamine (B_1_) and riboflavin (B_2_) suggest a basal state of high metabolic activity initiation. In our previous study, we have also assessed thiamine has a protective effect of biological agent under the fungal stress which is observable in whole genome expression profile (Silme et al. 2025). Thiamine pyrophosphate is a requisite cofactor for pyruvate dehydrogenase, linking glycolysis to the TCA cycle. The simultaneous upregulation of glutathione metabolism indicates that KB25 constitutively manages the oxidative stress inherent to saline environments, maintaining a redox balance essential for survival (Oren 1999). However, the *bacteria+fungus* interaction increased energy generation and proliferation. The dominant enrichment of ubiquinone and terpenoid-quinone biosynthesis in this group indicates that bacteria could give effort to supress pathogen fungus, likely to fuel the ATP-dependent production of secondary metabolites required for antagonism which can be correlated with the track of the remnant obtained from our MS analysis.

The metabolome of the *fungus alone* (in the context of the study’s comparative framework) was defined by profound perturbations in glycerophospholipid and arachidonic acid metabolism. While these pathways modulate signalling, in the context of microbial interactions, such extensive lipid variation often signals membrane stress or damage repair. If the fungus was exposed to bacterial metabolites (even prior to direct contact in the experimental design), these lipid shifts could represent a compensatory mechanism to maintain membrane fluidity and integrity against bacterial terpene-mediated disruption (Beccaccioli et al. 2019). The specific enrichment of flavone and flavonol biosynthesis might further reflect a fungal stress response, attempting to scavenge reactive oxygen species generated during competitive stress.

The docking simulation delineates a heterodimeric complex governed by high shape complementarity and a substantial interface area. The intimate steric fit between the bacterial ABC transporter and the plant glutamate receptor proteins suggests that this interaction is not merely a transient collisional encounter but a biologically specific assembly, likely following an induced fit mechanism. The dense network of hydrogen bonds and salt bridges identified by LigPlot+ highlights the role of electrostatic forces in conferring specificity to this interaction. The identification of Arg207(A) as a pivotal anchor is particularly significant. In PPIs, such residues are often classified as hot spots residues that contribute disproportionately to the binding free energy (Δ). The ability of Arg207(A) to bridge multiple residues (Asp51(B) and Lys298(B)) suggests it acts as an electrostatic clamp, locking the subunits in a precise orientation. Furthermore, the electrostatic interaction between Arg239(A) and Glu284(B), along with the peripheral stabilization by Asp4(A)/Glu2(A) and Gln75(B), likely contributes to electrostatic steering (Sinha and Smith-Gill 2002).

The stabilization of the central contact region by Tyr231(A) provides additional polar reinforcement, ensuring that the complex remains stable even if peripheral interactions are perturbed by solvent fluctuations. While electrostatics provide specificity, the stability of the complex appears to be thermodynamically driven by the hydrophobic effect. The Leucine-rich motif on the plant glutamate receptor proteins (Leu7, 10, 11, 15, 16, 18) forming a hydrophobic cluster with Val15, Ile16, and Pro111 on the bacterial ABC transporter is characteristic of high-affinity interfaces. The exclusion of water molecules from this non-polar core leads to a gain in solvent entropy, which is often the dominant favourable term in the Gibbs free energy equation for protein binding (Baldwin and Rose 2016).

In addition to metabolomics analyses, our findings have shown that KB25 protects plants against abiotic stress by regulating salt stress and also exhibits antagonistic properties against *P. infestans in vitro* conditions. Based on these findings, the close link between salt stress regulation and *P. infestans* inhibition in terms of mechanism of action is also supported by previous studies (Yan et al 2024; 2021). To finalize this study, we used the gene sequences of both proteins to model their interactions using docking analysis. We observed a significant interaction between osmotic betaine uptake transporter gene product of KB25 and glutamate receptor of tomato. This demonstrated the potential for substitution and suggested that this potential deficiency in the plant could be compensated for by KB25, which is attached to the rhizosphere and delivered by bacteria. Furthermore, we determined that terpene-derived compounds produced by KB25, whose presence we confirmed with genomic and metabolomic data, exert their inhibitory effect on *P. infestans* by binding to the RxLR effector, the virulence factor of *P. infestans* (Wang et al 2023; Resjö et al 2017). Since *P. infestans* lacks chitin in its cell wall, it can be suggested that an antagonistic effect with these terpene-derived compounds, rather than a possible enzymatic activity, is expected.

The KEGG pathway analysis indicated the existence of sporulenol as a terpene derivate. In the next step, we investigated the binding possibility of sporulenol to the effector proteins present in the disrupted cytoplasm, inactivating the RxLR effectors, and the pathogen loses its virulence. Because the pathogen cannot develop resistance to this ligand in a short time and this destructive effect gradually increases on pathogen cell.

Interestingly, we have found high structural similarity between the bacterial ABC transporter of KB25 and the plant glutamate receptor proteins of plant. We have also proved this similarity using ESPript and FoldScript servers (Gouet et al. 1999; Robert et al. 2025).

The genome-by-genome comparison output has also shown that the strain belongs to *T. aidigensis* confirmed with 16S rDNA analysis. But MLST analysis has indicated this strain as *Terribacillus halophilus* depending on allelic variation and housekeeping genes. Nevertheless, the ribosomal MLST analysis proved KB25 strain belongs to *T. aidigensis* (Online Resource 5). As suggested in our previous studies on bacterial identification, genome-by-genome comparison analysis or 16s ribosomal DNA sequencing alone is insufficient for taxonomic differentiation of halophilic bacteria (Bozkurt et al. 2025; Alatassi et al. 2025). Furthermore, ribosomal multi-locus sequence data will provide the most reliable results. We believe that in future studies, refraining from relying on a single piece of data for bacterial species identification will prevent future diagnostic errors and contribute to the accurate construction of taxonomic differentiation.

The pot experiment demonstrated that drenching with bacterial solution containing strain KB25 significantly alleviated salt stress in plants, as evidenced by reduced symptoms of salinity-induced inhibition compared to salt-stressed controls. This mitigation was associated with enhanced root formation and greater plant height in KB25-treated groups over 21 days post-treatment (Fig. 2), indicating a stress-regulatory role of the bacterium.

These observations align with established mechanisms of halotolerant or moderately halophilic plant growth-promoting rhizobacteria (PGPR), which promote plant resilience under saline conditions through multiple pathways. PGPR often enhance root architecture and biomass by producing phytohormones (e.g., IAA), facilitating nutrient uptake, and modulating osmotic balance via osmolyte accumulation or ion homeostasis. Inoculation frequently results in improved plant height, root length, and overall vigor in salt-stressed crops, as reported in various plant species treated with halotolerant strains (Radhakrishnan and Krishnasamy 2024). For example, halotolerant PGPR increase root development and shoot growth by regulating ABA signalling, reducing ethylene levels via ACC deaminase activity, and upregulating antioxidant defenses to counteract oxidative damage from NaCl stress (Giannelli et al. 2023). These previous findings are consistent with bacterial inoculation in our pot trials which mitigates salinity effects, leading to better growth parameters than uninoculated stressed conditions. These results highlight KB25’s potential as a bioinoculant for sustainable agriculture in saline soils, supporting crop productivity without chemical inputs. However, field trials, molecular analyses of stress-responsive genes, and identification of specific PGP traits in KB25 are needed to elucidate mechanisms and confirm efficacy in a large scale. Overall, this work underscores the role of stress-regulating bacteria in enhancing plant tolerance to salinity.

The metabolomics analysis indicated that co-cultivation induces substantial metabolic reprogramming, leading to altered production of specialized metabolites. In bacterial-fungal interactions, fungus often dominate the detectable metabolome, contributing key specialized compounds, while bacteria may modulate pathways or induce fungal responses. Enhanced production of protective metabolites under fungal challenge could contribute to antifungal antagonism and plant stress alleviation observed earlier, possibly via membrane disruption, nutrient competition, or oxidative stress induction in fungus (Masmoudi et al. 2021). These findings highlight co-culture metabolomics as a powerful strategy for uncovering inducible antifungal mechanisms in halotolerant bacteria like KB25. Comparatively, quinone biosynthesis and glycerophospholipids are shared across treatments, but interactions amplify cofactor (thiamine, CoA) and nucleotide pathways, absent or minor in monocultures, indicating KB25’s suppressive effect on the fungus via competitive resource allocation or elicitation of stress responses (Silme et al. 2025; Li et al. 2025b). *Fungus alone* lipid dominance may be attenuated in co-culture, favouring bacterial-driven energy pathways, aligning with KB25’s genomic osmoregulation and terpene features for antifungal antagonism (Ould Ouali et al. 2024). These interactions imply KB25 reprograms rhizosphere metabolism to mitigate salt stress (via osmolyte-linked cofactors) and suppress *P. infestans* via oxidative/energetic disruption, supporting its biocontrol potential. Future studies should validate specific metabolites’ roles in planta.

The metabolomic profiles across the three treatments *bacteria alone* (*T. aidingensis* KB25), *fungus alone* (*P. infestans*), and their co-culture highlight distinct metabolic signatures with shared motifs in cofactor biosynthesis, lipid remodelling, and energy metabolism, emphasizing KB25’s dual role in stress regulation and antifungal biocontrol.

In co-culture, ubiquinone/terpenoid-quinone biosynthesis (p=4.64×10^-4^) intensified antioxidants against interaction-induced stress, with vitamin K1 and α-tocotrienol. Thiamine (p=0.0038) and pantothenate/CoA (p=0.036) upregulated TCA/fatty acid metabolism, while purine (p=0.0024) and folate one-carbon pool (p=0.0042) supported nucleotide synthesis for repair. Glycerophospholipids (p=0.019), amino acid (p=0.0095), and benzoate degradation (p=0.042) converged on succinyl-CoA, fuelling energy demands. Shared quinone and glycerophospholipid motifs amplified cofactors and nucleotides in interactions, are absent in monocultures, suggesting KB25 suppresses *P. infestans* via resource competition or stress elicitation. This can be correlated with its antifungal activity against phytopathogens. Fungus lipid dominance attenuated, favouring bacterial energy shifts, linked to KB25’s osmoregulation genes and terpenes for antagonism. This reprograms rhizosphere metabolism to alleviate salt stress (osmolyte-cofactors) and *P. infestans* (oxidative disruption), enhancing biocontrol potential.

Integrated genomic-metabolomic analysis clarified the characteristics that confirmed our objective. This also demonstrated and guided the accuracy of our findings. It also confirmed the inhibitory role of sporulenol against pathogen and compensating of the glutamate receptor with the bacterial ABC-type proline/glycine betaine permease protein encoding in KB25 genome. Subsequently, integrated genomic-metabolomic analyses will be guide for further studies, due to its highly informative potential. We were also able to clearly see how the stress created by KB25 affects pathogen fungus metabolism.

The structural analysis of the bacterial ABC-type proline/glycine betaine permease protein and the glutamate receptor complex reveal a binding mechanism which shows the specific distribution of the residues identified in Fig. 6. The strong charge complementarity exemplified by Arg239(A), Glu284(B) and Glu2(A), Gln75(B) suggests that electrostatic steering plays a role in the initial recruitment of the subunits. The interaction between Tyr231(A) and Glu50(B) is of particular interest. Tyrosine residues at interfaces are frequently targets for phosphorylation. If Tyr231 is a phosphorylation site *in vivo*, the introduction of a phosphate group would introduce negative charge and steric bulk, likely disrupting the hydrogen bond with Glu50 and potentially abolishing the interaction. This suggests a plausible mechanism for the reversible regulation of this complex.

These structural protein-ligand docking analyses reveal the versatility of the sporulenol scaffold. It possesses the capacity to engage diverse protein targets through adaptable formation utilizing hydrophobic packing to inhibit pathogen virulence factors while exploiting polar surface interactions to potentially modulate host stress signalling. Previous studies have shown that the cell wall of *P. infestans* contains 1-6, 1-3 ß-glucans instead of chitin (Fry 2008); therefore, we investigated whether KB25 genome produce various glucanase enzymes. No gene region producing 1-6, 1-3 ß-glucanese enzymes was found in the genome data. It also became clearer that the suppressive effect on *P. infestans* is due to sporulenol production rather than cell wall degrading enzymes which was confirmed at both genomic and metabolomic analysis levels. Interestingly, we have proved that the content of RxLR protein are rich in terms of Arg-any amino acid-Leu-Arg motif and In our docking analysis we have found a strong interaction between sporulenol and Arg101 and 146, Leu14 which accordance with RxLR motif that could be linked to deformation of protein stability which is secreted proteins in *P. infestans* functioning to manipulate various pathogenic mechanisms involving manipulation of host immunity and pathogen fitness (Fry 2008).

## CONCLUSION

The comprehensive physiological, genomic, and metabolomic characterization of *T. aidigensis* strain KB25 highlights its strong potential as an effective biocontrol agent in saline agricultural environments. Its demonstrated antifungal activity against *P. infestans*, coupled with notable salt-stress tolerance, underscores its suitability for use in regions at which abiotic stress limits crop productivity. Genomic analyses revealing genes associated with secondary metabolite biosynthesis, along with metabolomic evidence of pathogen-inhibitory compounds, further validate the molecular basis of its antimicrobial action. In accordance with these findings the data not only confirm the strain’s capacity to suppress plant pathogens but also illustrate its adaptive mechanisms under osmotic stress, suggesting a dual role in both pathogen inhibition and enhancement of plant stress resilience. As such, strain KB25 represents a promising candidate for sustainable, biologically based disease management strategies in modern agriculture. Our findings have shown that sporulenol is a promising compound and therefore has the potential for use in the control of aerial mycelial fungal pathogens.

## ACKNOWLEDGEMENT

This study dedicated to memory of late Prof. Dr. Hans-Heinrich Hoppe who was one of Ph.D. advisors’ Dr. Ömür Baysal from Georg August University, Göttingen, Germany.

## COMPLIANCE WITH ETHICAL STANDARDS

This article does not contain any studies with human participants.

## AUTHOR CONTRIBUTIONS

GKA: Formal analysis; Investigation; Methodology; Resources; Writing - original draft. ÖB: Conceptualization, Data curation; Formal analysis; Funding acquisition; Investigation; Methodology; Project administration; Resources; Software; Supervision; Validation; Visualization; Writing - original draft; Writing - review & editing. RSS: Formal analysis; Funding acquisition; Methodology; Project administration; Writing - original draft; Writing - review & editing. ÜÇ: Formal analysis; Methodology; Visualization.

## DATA AVAILABILITY STATEMENT

The raw sequence data of this study are openly available in the NCBI database (Acc. No: PRJNA1375849 and PX658357.1). The data supporting the findings of this study are presented in the supplementary material of this article.

## SUPPLEMENTARY MATERIALS

(1) TOF/MS running gradient conditions for metabolomics analysis.
(2) Temperature-dependent changes in antibiotic susceptibility variations of KB25.
(3) Genome annotation and mapping of whole genome of KB25.
(4) The pie chart depicts the number of gene locations encoding proteins of KB25 genome.
(5) rST analysis output of KB25.
(6) The antiSMASH results of KB25 genome.
(7) Number of *T. aidingensis* KB25 genes related to various molecular functions and cellular components.
(8) Metabolomics results.
(9) TM-align and Foldseek results.
(10) Protein-protein docking output.
(11) Protein-ligand AutoDock output related to RxLR effector peptide protein of putative (*Phytophthora infestans* T30-4) and ligand sporulenol.
(12) Protein-ligand AutoDock output of glutamate receptor and ligand sporulenol.

